# Supercharged Cellulases Show Reduced Non-Productive Binding, But Enhanced Activity, on Pretreated Lignocellulosic Biomass

**DOI:** 10.1101/2021.10.17.464688

**Authors:** Bhargava Nemmaru, Jenna Douglass, John M Yarbrough, Antonio De Chellis, Srivatsan Shankar, Alina Thokkadam, Allan Wang, Shishir P. S. Chundawat

## Abstract

Non-productive adsorption of cellulolytic enzymes to various plant cell wall components, such as lignin and cellulose, necessitates high enzyme loadings to achieve efficient conversion of pretreated lignocellulosic biomass to fermentable sugars. Carbohydrate-binding modules (CBMs), appended to various catalytic domains (CDs), promote lignocellulose deconstruction by increasing targeted substrate-bound CD concentration but often at the cost of increased non-productive enzyme binding. Here, we demonstrate how a computational protein design strategy can be applied to a model endocellulase enzyme (Cel5A) from *Thermobifida fusca* to allow fine-tuning its CBM surface charge, which led to increased hydrolytic activity towards pretreated lignocellulosic biomass (e.g., corn stover) by up to ∼330% versus the wild-type Cel5A control. We established that the mechanistic basis for this improvement arises from reduced non-productive binding of supercharged Cel5A mutants to cell wall components such as crystalline cellulose (up to 1.7-fold) and lignin (up to 1.8-fold). Interestingly, supercharged Cel5A mutants that showed improved activity on various forms of pretreated corn stover showed increased reversible binding to lignin (up to 2.2-fold) while showing no change in overall thermal stability remarkably. In general, negative supercharging led to increased hydrolytic activity towards both pretreated lignocellulosic biomass and crystalline cellulose whereas positive supercharging led to a reduction of hydrolytic activity. Overall, selective supercharging of protein surfaces was shown to be an effective strategy for improving hydrolytic performance of cellulolytic enzymes for saccharification of real-world pretreated lignocellulosic biomass substrates. Future work should address the implications of supercharging cellulases from various families on inter-enzyme interactions and synergism.

## INTRODUCTION

The future circular economy is based on conversion of wastes from a variety of streams to useful products that are currently produced from fossil fuels^1,2^. Bioethanol is one such product that can be potentially produced from lignocellulosic biomass such as agricultural residues (e.g., corn stover, wheat/rice straws, sugarcane bagasse) and forest residues (e.g., wood chips)^3^. The versatility of available biomass sources and the variety of bioproducts that can be generated, lends itself to development of customized conversion strategies tailor-made for various feedstocks in an integrated biorefinery^4,5^. One conversion strategy that has received significant attention is the enzymatic conversion of cellulose and hemicellulose to C6/C5 based mixed sugar streams^3^, while employing tailored valorization strategies for extracted lignin based on the pretreatment strategy^6,7^. These sugars can be converted to a variety of platform chemicals such as ethanol, organic acids, or polymer-precursors in an integrated biorefinery^8^.

Various techno-economic analyses have been performed to assess the feasibility of producing bioethanol in a cost-effective and sustainable manner from biomass^9,10^. These studies have highlighted the role of prohibitively high enzyme costs towards commercialization of biofuels^11^. Hence, there is a need to develop enzyme engineering strategies to improve the overall conversion of lignocellulosic biomass to reducing sugars, while reducing biomass recalcitrance via thermochemical pretreatment^12,13^. Non-productive enzyme binding to lignin and cellulose along with limited enzyme accessibility to the substrate are considered the key deterrents of enzyme activity towards pretreated biomass substrates^14–17^. As a result, pretreatment efforts have focused on extraction of lignin for valorization while also improving overall enzyme accessibility to the residual polysaccharides^18,19^.

However, most pretreatment technologies (e.g., dilute acid, extractive ammonia, alkaline, deacetylation and mechanical refining or DMR) only extract lignin partially, leaving behind residual lignin that can still deactivate or inhibit enzymes^20,21^. Lignin has been shown to deactivate cellulases through various mechanisms, the most significant of which involves protein conformational changes upon adsorption to lignin driven via hydrophobic interactions^22–24^. Broadly speaking, the three strategies that have been employed to reduce cellulase non-productive binding to lignin include: (i) addition of sacrificial proteins such as BSA^25^ or soy protein,^26^ (ii) inclusion of negatively charged groups such as acetyl groups on the surface of enzymes via chemical conjugation,^27^ and (iii) enzyme surface supercharging via computational re-design^28,29^. Although the first two strategies have been shown to reduce lignin inhibition, they require addition of an additional reagent (BSA or soy protein) or treatment procedure (acetylation), which increases the operating or capital cost of the bioconversion process. On the other hand, enzyme supercharging is an inexpensive method of genetically engineering enzymes to alter their surface electrostatic properties^30,31^.

Protein supercharging has been used to accomplish a variety of useful applications including but not limited to macromolecule or drug delivery into mammalian cells^32^, DNA detection and methylation analysis^33^, complex coacervation with polyelectrolytes^34^, self-assembly into organized structures^35^ such as protein nanocages^36^ and Matryoshka-type structures^37^ and encapsulation of cargo proteins into such higher-order structures^38^. Previously, we have utilized a supercharging strategy based on Rosetta^39,40^ and FoldIt standalone interface^41^ for engineering green fluorescent protein (GFP)^42^ and CelE (from *Ruminiclostridium thermocellum*)^29^. We found that net negative charge was correlated weakly with reduced lignin binding capacity for GFP supercharged mutants, whereas the charge density was not found to have a clear impact on lignin binding capacity^42^. In our follow-up study^29^, a cellulase catalytic domain CelE was fused with CBM3a and both domains were individually negatively supercharged. Negatively supercharged CBM3a designs showed relatively improved hydrolysis yields on model amorphous cellulose in the presence of lignin, compared to the wild-type enzyme. However, all tested designs showed reduced absolute activity than wild-type controls on amorphous cellulose substrates (and with no data reported on pretreated lignocellulosic biomass) which was hypothesized to be due to reduced binding to cellulose induced by electrostatic repulsions.

Although these studies show proof-of-concept for the potential beneficial impact of cellulase negative supercharging on biomass hydrolysis, there still remain multiple unanswered mechanistic questions to fully leverage the potential of enzyme supercharging for lignocellulosic biomass hydrolysis. Broadly speaking, there are four unanswered questions: (i) how does negative supercharging impact enzyme kinetics on pretreated biomass substrates? (ii) is there an optimal net negative charge range which results in maximum biomass hydrolysis yields? (iii) how can biomass hydrolysis performance be rationalized by understanding the relative impact of non-productive enzyme binding to cellulose and/or lignin on overall biomass hydrolysis yields? (iv) finally, could enhanced thermal stability also have a significant contribution towards improved activity? Here, we sought to address these questions in greater detail, using a model endocellulase enzyme Cel5A from *Thermobifida fusca (T. fusca)*, which has been well-characterized in our lab previously^43^. *T. fusca* is a thermophilic microbe that secretes cellulase enzymes belonging primarily to glycosyl hydrolase (GH) families 5, 6, 9, and 48, with most cellulase CDs tethered to a type-A CBM2a^44^. Testing the protein supercharging strategy on a model Cel5A enzyme and it’s CBM2a from this cellulolytic enzyme system will also allow for extension of these design principles to other enzymes, potentially leading to a supercharged cellulase mixture with superior performance.

More specifically, we computationally designed a library of eight CBM2a designs spanning a net charge range of -14 to +6. These CBM2a designs were fused with the Cel5A catalytic domain and green fluorescent protein (GFP) separately, to study the hydrolysis activity and binding behavior of the constructs on a variety of substrates, respectively. Firstly, we characterized the hydrolysis yields of CBM2a-Cel5A fusion constructs at various reaction times (2-24 h) on ammonia fiber expansion (AFEX) and extractive ammonia (EA) pretreated corn stover substrates. To deconvolute the contributions from various factors such as improved thermal stability, non-productive binding to cellulose and lignin, we performed thermal shift assays and follow-on hydrolysis assays towards isolated crystalline cellulose substrates in the presence and absence of lignin. Moreover, we performed binding assays to study the binding of GFP-CBM2a fusion constructs to cellulose and lignin using previously established QCM-D assay procedures^17,42^. Overall, our results identify the mechanism by which CBM supercharging can enhance activity and highlight the utility of cellulolytic enzyme supercharging to improve pretreated biomass hydrolysis yields in a biorefinery relevant context for the first time.

## EXPERIMENTAL SECTION

### Reagents

AFEX and EA pretreated corn stover were prepared and provided in kind by Dr. Rebecca Ong’s lab (Michigan Technological University, Houghton) and Bruce Dale’s lab (Michigan State University, East Lansing), according to previously established protocols^45–47^. Avicel (PH 101, Sigma-Aldrich, St Louis) was used to prepare cellulose-III allomorph with the following pretreatment conditions (90 °C, 6:1 anhydrous liquid ammonia to cellulose loading, and 30 minutes of total residence time) and phosphoric acid swollen cellulose (PASC) as described previously^48^. Sarvada Chipkar from the Ong lab kindly prepared and provided cellulose-III used in this study. Lignin extracted from corn stover was prepared using the organosolv extraction process^49^ and kindly provided by Stuart Black of the National Renewable Energy Laboratory (NREL). All other chemicals and analytical reagents were procured either from Fisher Scientific or Sigma Aldrich, or as noted in the relevant experimental section.

### Mutant energy scoring using Rosetta

Creation of computational designs carrying a certain net charge necessitated computing the change in energy scores upon mutation of a native amino acid residue to either a positively charged (K, R) or a negatively charged residue (D, E). These mutations were scored using Rosetta. The wild-type protein PDB file is obtained either via homology modeling using Rosetta CM^50^ or via the protein data bank^51^. PyMOL (Schrodinger) was used to generate the desired mutation in amino acid sequence of a given protein and exported as a PDB file that represents the mutated protein. Customized scripts were developed in Rosetta to perform fast relax^52^ of any input PDB file. PDB files of both the wild-type and mutated proteins were relaxed separately using ten fast relax operations at a time. Each round of energy minimization enabled by ten fast relax operations was repeated until the Rosetta energy score of protein equilibrated and did not vary by more than 0.1 Rosetta Energy Units (REU) between one round of energy minimization (comprising of 10 fast relaxes) to another. The mutation energy score for a given mutation was calculated by measuring the difference between Rosetta energy scores of the wild-type protein and the mutant after energy minimization.

### Plasmid generation, protein expression and purification

*Thermobifida fusca* native Cel5A^53,54^ (UniprotKB - Q01786) gene was cloned into pET28a(+) (Novagen) and was kindly provided by Nathan Kruer-Zerhusen (from late Prof. David Wilson’s lab at Cornell University). An N-terminal 8X His tag was inserted and the native signal peptide removed from the original gene construct. The gene was then cloned into our in-house expression vector pEC to optimize protein expression yields as described previously^55,56^. The plasmid maps for pEC-CBM2a-Cel5A and pEC-GFP-CBM2a are provided in **SI Appendix Figures S1** and **S2** respectively. The full nucleotide sequences with color coding for each gene segment are reported in a separate supplemental file (see **SI Appendix Sequences**). CBM2a mutant designs were ordered from Integrated DNA Technologies, Inc (IDT) as custom-synthesized gBlocks. These CBM2a design gBlocks (see **SI Appendix Table T1** for the CBM2a design nucleotide sequences) were then swapped with wild-type CBM2a to generate mutant CBM2a-Cel5A fusion constructs using standard sequence and ligation independent cloning (SLIC) protocols. A similar approach was used to insert CBM2a designs into previously reported pEC-GFP-CBM vector^56^. Molecular cloning for *Thermobifida fusca* β-glucosidase (UniprotKB - Q9LAV5) gene These colonies were then inoculated in LB medium and grown overnight to prepare 20% glycerol stocks for long-term storage at -80 °C. These glycerol stocks were then used to inoculate 25 ml of LB media with 50 µg/ml kanamycin and incubated at 37 °C, 200 rpm for 16 hours. These overnight cultures were then transferred to 500 ml auto-induction medium (TB+G)^57^ and incubated at 37 °C, 200 rpm for 6 hours to allow optical density to reach the exponential regime. Protein expression was then induced by reducing the temperature to 25 °C for 24 hours at 200 rpm. Cell pellets were then harvested using Beckman Coulter centrifuge and JA-14 rotor by spinning the liquid cultures in 250 ml plastic bottles at 30,100 g for 10 minutes at 4 °C. All the cell culturing experiments were performed using an Eppendorf Innova™ incubator shaker. Cell pellets were lysed using 15 ml cell lysis buffer (20 mM phosphate buffer, 500 mM NaCl, 20% (v/v) glycerol, pH 7.4), 0.5 mM Benzamidine (Calbiochem 199001), 200 µl protease inhibitor cocktail (1 µM E-64 (Sigma Aldrich E3132), 15 µl lysozyme (Sigma Aldrich, USA) and 1 mM EDTA (Fisher Scientific BP1201)) for every 3 g wet cell pellet. The cell lysis mixture was sonicated using Misonix™ sonicator 3000 for 5 minutes of total process time at 4.5 output level and specified pulse settings to avoid sample overheating (pulse-on time: 10 seconds and pulse-off time: 30 seconds). An Eppendorf centrifuge (5810R) with F-34-6-28 rotor was then used to separate the cell lysis extract from insoluble cellular debris at 15,500 g, 4 °C for 45 minutes. Immobilized metal affinity chromatography (IMAC) using His-Trap FF Ni^+2^-NTA column (GE Healthcare) attached to BioRad™ NGC system, was then performed to purify the his-tagged proteins of interest from the background of cell lysate proteins. Briefly, there were three steps involved during IMAC purification: 1. equilibration of column in buffer A (100 mM MOPS, 500 mM NaCl, 10 mM Imidazole, pH 7.4) at 5 ml/min for 5 column volumes, 2. soluble cell lysate loading at 2 ml/min, and 3. His-tagged protein elution using buffer B (100 mM MOPS, 500 mM NaCl, 500 mM Imidazole, pH 7.4). The purity of eluted proteins was validated using SDS-PAGE before buffer exchange into 10 mM sodium acetate (pH 5.5) buffer for long-term storage after flash freezing at -80 °C and/or follow-on activity characterization.

### Pretreated lignocellulosic biomass hydrolysis assays

AFEX and EA corn stover (milled to 0.5 mm) were suspended in deionized water to obtain slurries of 25 g/l total solids concentration. All biomass hydrolysis assays were performed in 0.2-ml round-bottomed microplates (PlateOne™), with at least 4 replicates for each reaction condition. Reactions quenched at different time points (2, 6 and 24 hours) were performed in different microplates. Each reaction was composed of 80 μl biomass slurry (25 g/l), 20 μl sodium acetate buffer (0.5 M), 50 μl cellulase enzyme (at appropriate concentration), 25 μl β-glucosidase (at appropriate concentration), and 25 μl of deionized water to make up the total reaction volume to 200 μl. For reaction blanks, the enzyme solutions were replaced with deionized water while biomass slurry and buffer volumes remained the same. The cellulase enzyme loading was maintained at 120 nmol per gram biomass substrate and the β-glucosidase enzyme loading was maintained at 12 nmol per gram biomass substrate (leading to 10% of cellulase enzyme concentration). Upon addition of all the requisite reaction components, the microplates were covered with a plate mat, sealed with packaging tape, and incubated at 60 °C for the specified time duration (2, 6 or 24 hours) with end-over-end mixing at 5 rpm in a VWR hybridization oven. Upon reaction completion, the microplates were centrifuged at 3900 rpm for 10 minutes at 4 °C to separate the soluble supernatant (comprised of soluble reducing sugars) from insoluble biomass substrate. The supernatants were then recovered and dinitrosalicylic acid (DNS) assays were performed as previously described to estimate total soluble reducing sugars^43^. This data was fitted to a two-parameter kinetic model that was previously deployed to study reaction kinetics of *T. fusca* cellulases on biomass substrates^59^. Origin software was used to perform the curve fitting analysis and obtain the pseudo-kinetic time-dependent parameters ‘A’ and ‘b’ which represent the net activity of bound enzyme and the time-dependent ability of enzyme to overcome recalcitrance, respectively. An increase in b might indicate the ability of enzyme to sample new substrate sites as reaction progresses, thereby reducing substrate recalcitrance.

### Cellulose hydrolysis assays and lignin inhibition assays

The cellulose hydrolysis assays were performed in a similar manner as biomass hydrolysis assays, except for the reaction composition. Avicel PH101 derived cellulose-I and cellulose-III were suspended in deionized water to form slurries of 100 g/l total solids concentration. A 0.2-ml round-bottomed microplate (PlateOne™) was used for each discrete reaction timepoint (2, 6 and 24 hours) and each reaction was performed with at least 4 replicates. Each reaction was composed of 40 μl cellulose slurry (100 g/l), 20 μl sodium acetate buffer (0.5 M, pH 5.5), 50 μl cellulase enzyme (at appropriate concentration), 25 μl β-glucosidase (at appropriate concentration) and 65 μl of deionized water to make up the total reaction volume to 200 μl. The cellulase enzyme loading was maintained at 120 nmol per gram biomass substrate and the β-glucosidase enzyme loading was maintained at 12 nmol per gram biomass substrate (leading to 10% of cellulase enzyme concentration). Upon reaction completion, supernatants were removed, and DNS assays were performed as described in the previous section on biomass hydrolysis assays. On the other hand, the reaction mixture for lignin inhibition assays was composed of 20 μl cellulose slurry (100 g/l), 40 ul lignin slurry (20 g/l), 20 μl sodium acetate buffer (0.5 M, pH 5.5), 50 μl cellulase enzyme (at appropriate concentration), 25 μl β-glucosidase (at appropriate concentration) and 65 μl of deionized water to make up the total reaction volume to 200 μl. The enzyme loadings and all the follow-on steps were conducted in a similar manner to cellulose hydrolysis assays. 24 hours was used as the preferred reaction time for lignin inhibition assays, owing to the prevalence of lignin and cellulose non-productive binding at longer reaction times.

### Quartz crystal microbalance with dissipation (QCM-D) based binding assays

Preparation of cellulose and lignin films for characterization of GFP-CBM binding, was performed as described elsewhere^17,60^. Quartz sensors functionalized with nanocrystalline cellulose or lignin were mounted on the sensor holder of QSense E4 instrument and equilibrated with buffer (50 mM sodium acetate, pH 5.5 with 100 mM NaCl) for 10 minutes at a flow rate of 100 μl/min using a peristaltic pump. The films were left to swell in buffer overnight and the films were considered stable if the third harmonic reached a stable baseline after overnight incubation. GFP-CBM2a protein stocks were then diluted to a concentration of 2.5 μM using 50 mM sodium acetate (pH 5.5) and flown over the sensors at a flow rate of 100 μl/min for 10 – 15 minutes until the system reached saturation, as observed by the third harmonic. The system was then allowed to equilibrate for at least 30 minutes and protein unbinding was then tracked by flowing buffer (50 mM sodium acetate, pH 5.5 with 100 mM NaCl) over the sensors at a flow rate of 100 μl/min for at least 30 minutes. Data analysis for QCM-D traces was performed as described previously^17^. However, for lignin, binding was observed to be mostly irreversible^61^ and hence, only the maximum number of binding sites and percent irreversible protein bound, calculated based on the maximum number of binding sites and the amount of protein bound towards the end of unbinding regime.

### Cellulase thermal shift assay

The protocol for thermal shift assays was similar to that reported previously^29^. Briefly, 5 µl 200X SYPRO reagent, 5 µl 0.5 M sodium acetate buffer (pH 5.5), enzyme dilution to make up an effective concentration of 5 µM and deionized water to make up the total volume to 50 µl were added to MicroAmp™ EnduraPlate™ 96-well clear microplate (Applied Biosystems™). QuantStudio3 (Applied Biosystems™) was then used to measure the fluorescence using the channel allocated to FAM dye (excitation: 470 nm, emission: 520 nm) under a temperature ramp from 25 °C to 99 °C at a rate of 0.04 °C per second. The melting curves obtained were then analyzed using an open-source tool called SimpleDSFViewer^58^.

## RESULTS AND DISCUSSION

### Computational protein design strategy

A homology model was constructed for the target CBM2a protein using Rosetta CM tool^50^ based on templates from CBM family 2a with at least 50% sequence identity. Surface residues were then identified using an appropriate residue selector in Rosetta. Previous studies have shown that 10% of the total amino acid sequence length of globular proteins can be mutated using the supercharging strategy, while still allowing the proteins to fold properly^31^. Given that CBM2a is 100 amino acids long and has a net charge of -4, we sought to generate designs that spanned a net charge range of -14 to +6 using 10 mutations of polar uncharged amino acid residues. Overall, 31 polar uncharged amino acid residues were identified on the protein surface and these residues were scored individually for mutations to lysine (K), arginine (R), aspartic acid (D) and glutamic acid (E).

The mutation energy scores were then averaged for any given position and surface polar uncharged residues were sorted based on these average mutation energy scores. From the original pool of 31 polar uncharged residues, three categories of residues were considered immutable due to their potential implications for protein folding or interaction with cellulose as follows: 1. residues within 10 Å distance from evolutionarily conserved planar aromatic residues^62^ essential for CBM function, 2. residues on the CBM binding face^63^, and 3. residues with a positive average mutation energy score (predicting structural instability upon mutation). Upon exclusion of these three categories of residues, 11 mutable polar uncharged residues were identified and sorted into two spatially distinct clusters and sorted based on their mutation energy scores from highest to lowest. The individual and average mutation energy scores of mutable residues are reported in **SI Appendix Table T2**. Eight designs were then generated to have net charges of -14 (D1), -12 (D2), -10 (D3), -8 (D4), -6 (D5), -2 (D6), +2 (D7) and +6 (D8) as shown in **Figure 1**. The mutations used to generate each design, are reported in **SI Appendix Table T3** whereas the full amino acid sequence for wild-type CBM2a with these mutable residues highlighted in red font, are reported in a separate supplemental file (see **SI Appendix Sequences**). For instance, for generation of design D5 (−6 net charge) from wild-type (−4 net charge), N39 and T49 positions were picked for mutation since they were higher up in the sorted list of residues in their individual clusters as shown in **SI Appendix Table T3**.

**Figure 1.**
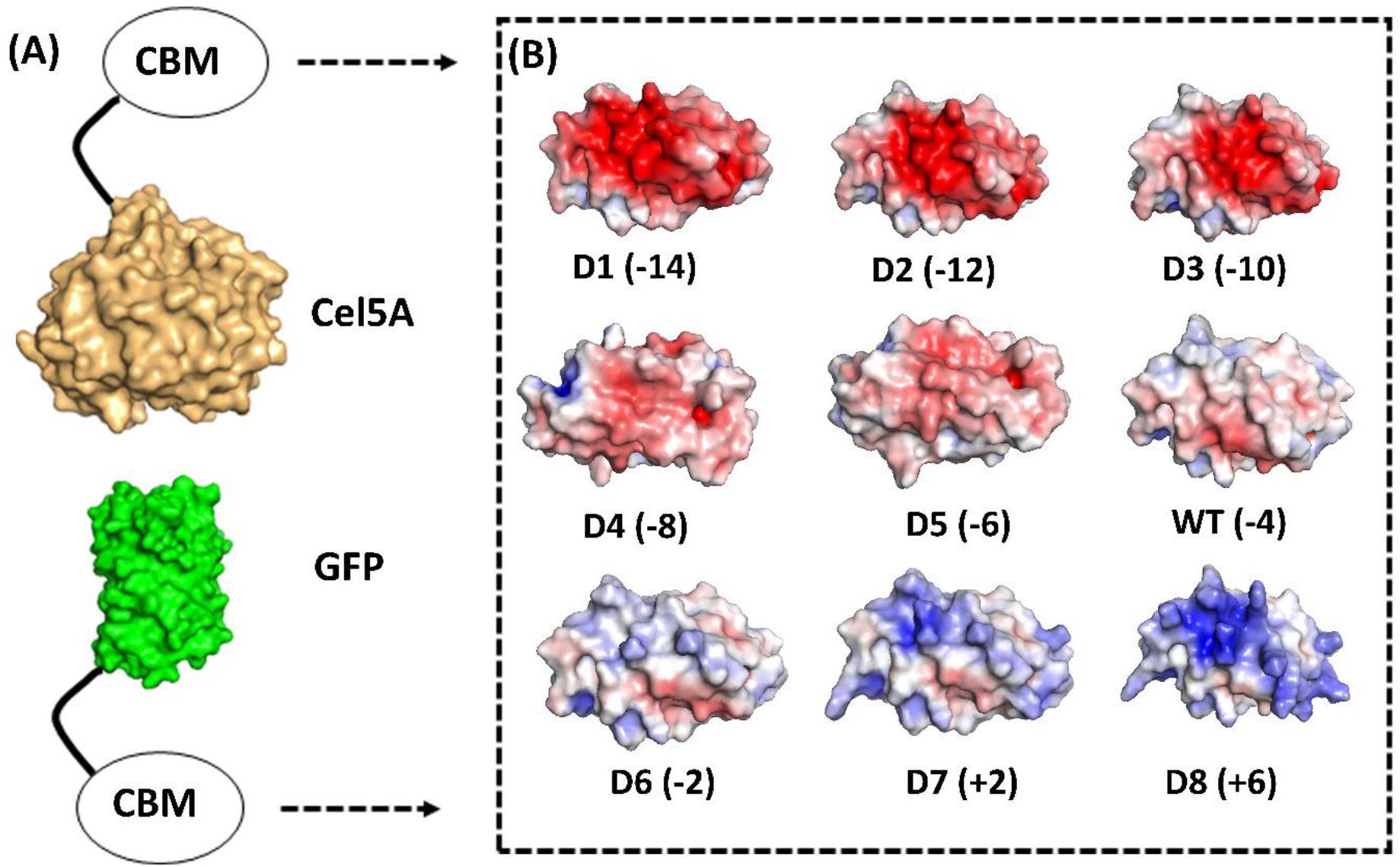
Computational design of supercharged CBM2a mutants and generation of fusion protein constructs. Rosetta was used to identify amino acids on the surface of CBM2a wild-type protein which are amenable to positively charged (K, R) or negatively charged (D, E) amino acid mutations to achieve a target net charge spanning the -14 to +6 range. (A) CBM2a designs were fused with Cel5A and GFP separately. (B) Electrostatic potential maps of the 8 CBM2a designs and their wild-type (represented as WT) are generated using APBS Electrostatics Tool in PyMOL. The name of each construct (D1 – D8 and WT) is followed by the net charge of each design in parenthesis.

### Negatively supercharged cellulases show improved hydrolysis yields on pretreated corn stover

All the CBM2a-Cel5A and GFP-CBM2a designs were cloned, expressed, and purified as described in the experimental procedures section. Here, we tested the hydrolytic activity of our supercharged CBM2a-Cel5A designs against two well-characterized pretreated biomass substrates, namely AFEX and EA corn stover. These two substrates were chosen with the intent of screening enzyme activities against substrates with differing lignin content. Hydrolysis yields were measured in terms of percent conversion (based on theoretical reducing sugar yield from both cellulose and hemicellulose) at three different time points (2 hr, 6 hr and 24 hr) at 120 nmol enzyme loading per gram substrate, resulting in reaction progress curves shown in **Figure 2A and 2C**, for AFEX and EA corn stover respectively.

**Figure 2.**
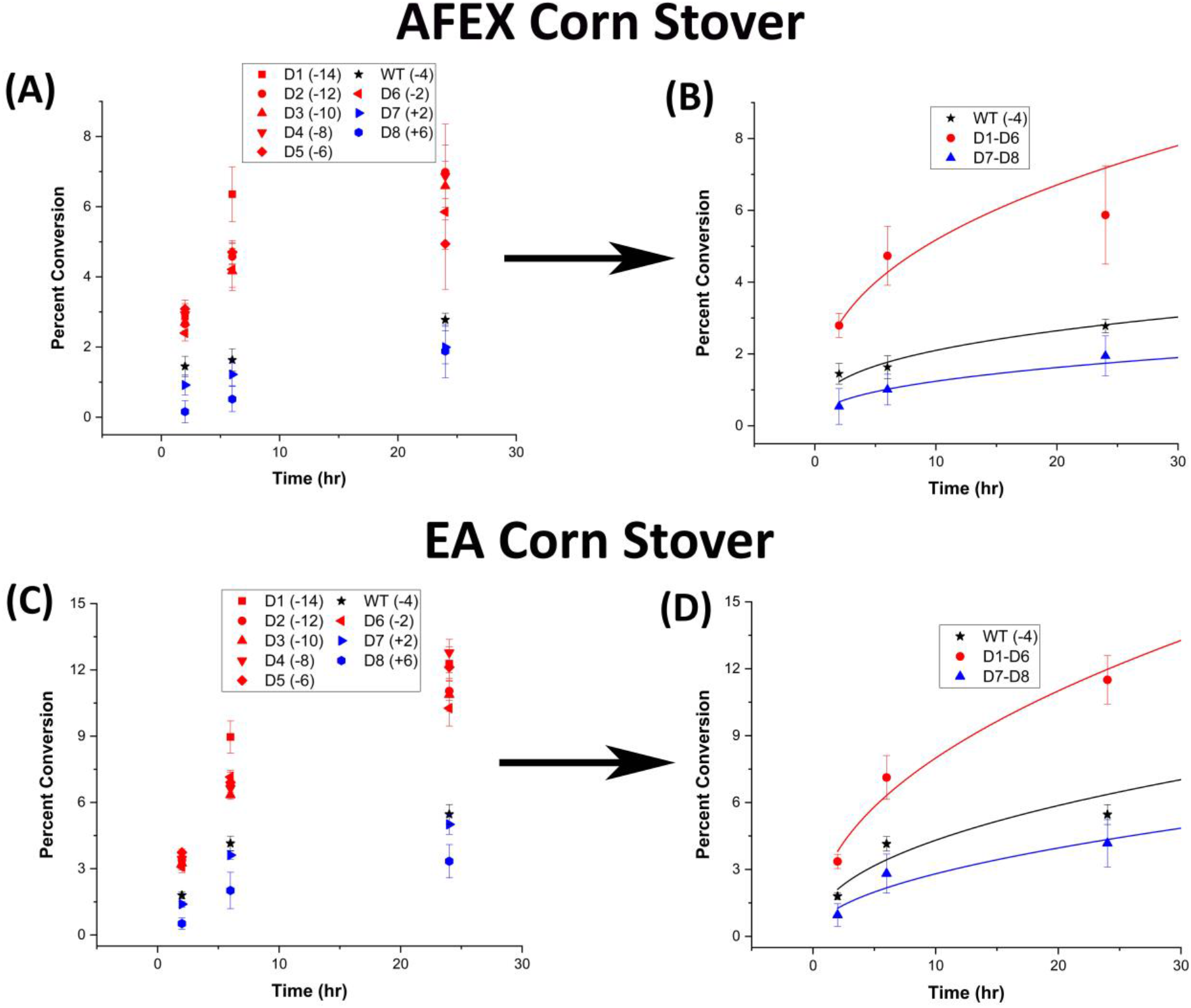
Negatively supercharged enzymes show improved hydrolysis yields on pretreated biomass substrates. 80 μL of 25 g/L biomass substrate was hydrolyzed using an enzyme loading of 120 nmol CBM2a Cel5A fusion enzyme per gram biomass substrate with 12 nmol β-glucosidase enzyme (10% of cellulase loading) per gram biomass substrate for reaction times of 2hrs, 6hrs, and 24hrs. The solubilized reducing sugar concentrations in the supernatant after hydrolysis were determined by the DNS assay. Percent conversion of biomass to glucose equivalents is reported on the Y-axis. (A) Percent conversion as a function of time (2 hr, 6 hr and 24 hr) for the hydrolysis of AFEX Corn Stover by D1 – D8 CBM2a Cel5A and WT CBM2a Cel5A. Error bars represent standard deviation from the mean, based on at least 4 replicates. (B) Based on the data reported in (A), CBM2a Cel5A fusion constructs with negatively charged CBMs (D1 – D6) were grouped together and average hydrolysis yields were obtained for the group, with the error bars representing standard deviation from the mean. Similarly, CBM2a-Cel5A fusion constructs with positively charged CBMs (D7 and D8) were grouped together and average hydrolysis yields were obtained. Trend curves have been added to represent the kinetic profiles of the hydrolysis reaction. (C) Percent conversion as a function of time (2 hr, 6 hr and 24 hr) for the hydrolysis of EA Corn Stover by D1 – D8 CBM2a Cel5A and WT CBM2a Cel5A. Error bars represent standard deviation from the mean, based on at least 4 replicates. (D) Based on the data reported in (C), CBM2a Cel5A fusion constructs with negatively charged CBMs (D1 – D6) were grouped together and average hydrolysis yields were obtained for the group, with the error bars representing standard deviation from the mean. Similarly, CBM2a-Cel5A fusion constructs with positively charged CBMs (D7 and D8) were grouped together and average hydrolysis yields were obtained. Trend curves have been added to represent the kinetic profiles of the hydrolysis reaction.

As shown in **Figure 2A and 2C**, all CBM2a-Cel5A fusion constructs with a net negative charge on the CBM (D1 – D6) showed higher activity than the wild-type enzyme whereas those with positively charged CBM (D7 and D8) showed reduced activity towards AFEX and EA corn stover at all the reaction times considered. The reaction kinetics data was then fit to a two-parameter model as described previously^59^ and the parameter fits are reported in **SI Appendix Table T4**. Parameter ‘A’ represents the net activity of bound enzyme whereas the parameter ‘b’ represents the enzymes ability to reduce biomass recalcitrance over time. For AFEX corn stover, the negatively charged (D1 – D6) CBM2a-Cel5A fusion constructs showed ∼ 2.1 to 3.3-fold improvement in ‘A’ compared to the wild-type, with D1 showing the most improvement. However, these constructs exhibited a value of ‘b’ comparable to the wild-type. On the other hand, for EA corn stover, the negatively charged (D1 – D6) CBM2a-Cel5A fusion constructs showed ∼ 1.5 to 2.2-fold improvement in ‘A’ whereas ‘b’ remained comparable to the wild-type. The positively charged (D7 and D8) CBM2a-Cel5A constructs showed a reduction in ‘A’ regardless of the biomass substrate, with D8 consistently performing the worst, with ∼ 4.8 and ∼ 3.3-fold reduction towards AFEX and EA corn stover respectively. These results indicate that negative supercharging might improve the net activity of bound enzyme (indicated by A) towards alkaline pretreated biomass substrates whereas the ability to overcome recalcitrance by identifying new substrate sites (indicated by b) might be still limited.

Student’s T-tests were performed between hydrolysis yields of mutants within the negative net charge (D1 – D6) and positive net charge (D7 and D8) groups which did not reveal statistically significant differences (see **SI Appendix Table T5** for Student’s T-test results). Hence, the mutants in these groups could be classified together and mean hydrolysis yields were obtained for each sub-group (D1 – D6 vs D7 – D8), to further understand the high-level impact of positive vs negative net charge of supercharged cellulases on biomass hydrolysis. These results are represented in **Figure 2B and 2D** for AFEX and EA corn stover, respectively.

Towards AFEX corn stover (**Figure 2B**), the negatively charged group of enzymes showed up to 2 to 3-fold improvement over the wild-type enzyme, with the maximum fold-improvement being seen at the earlier time points (i.e., 6 hours). However, the positively charged group showed ∼ 1.5 to 2.5-fold reduction in hydrolysis yields. A similar trend was observed in the case of EA corn stover although the improvements for positively charged sub-group and reductions for the negatively charged sub-group were relatively modest. The kinetic parameter fits for these groups are also listed in **SI Appendix Table T4** and the negatively charged group shows ∼ 2.2 and 1.8-fold improvement in ‘A’ model parameter compared to the wild-type enzyme, towards AFEX and EA corn stover, respectively.

This improvement in biomass hydrolysis yields for the negatively charged cellulase group could arise from a combination of various factors: (i) an increase in specific activity towards cellulose due to reduced non-productive binding to cellulose, (ii) reduced non-productive binding to lignin, and (iii) improved thermal stability. Here, we tested each of these hypotheses systematically. Prior to studying the binding of supercharged CBM2a designs to cellulose and lignin, we sought to test the activities of enzyme designs towards cellulose-I and cellulose-III in the presence or absence of lignin. The choice of cellulose-I and cellulose-III is driven by these allomorphs being the predominant forms in AFEX and EA corn stover, respectively.

### Negatively supercharged cellulases also show improved hydrolysis yields on crystalline cellulose allomorph substrates

Assays were conducted to quantify the impact of CBM supercharging on crystalline cellulose hydrolysis. Hydrolysis yields were measured in terms of percent conversion at three different time points (2 hr, 6 hr and 24 hr) at 120 nmol enzyme per gram substrate loading, resulting in reaction progress curves shown in **Figure 3A** and **Figure 3C** for cellulose-I and cellulose-III, respectively. Cellulose-I and cellulose-III were chosen as target substrates since AFEX and EA corn stover are comprised of these two cellulose allomorphs, respectively. Similar to the behavior observed with pretreated biomass, Cel5A fusion constructs tethered with negatively charged CBM2a (D1-D6) showed improved activity towards both cellulose-I and cellulose-III whereas constructs with positively charged CBM2a (D7 – D8) designs showed reduced activity.

**Figure 3:**
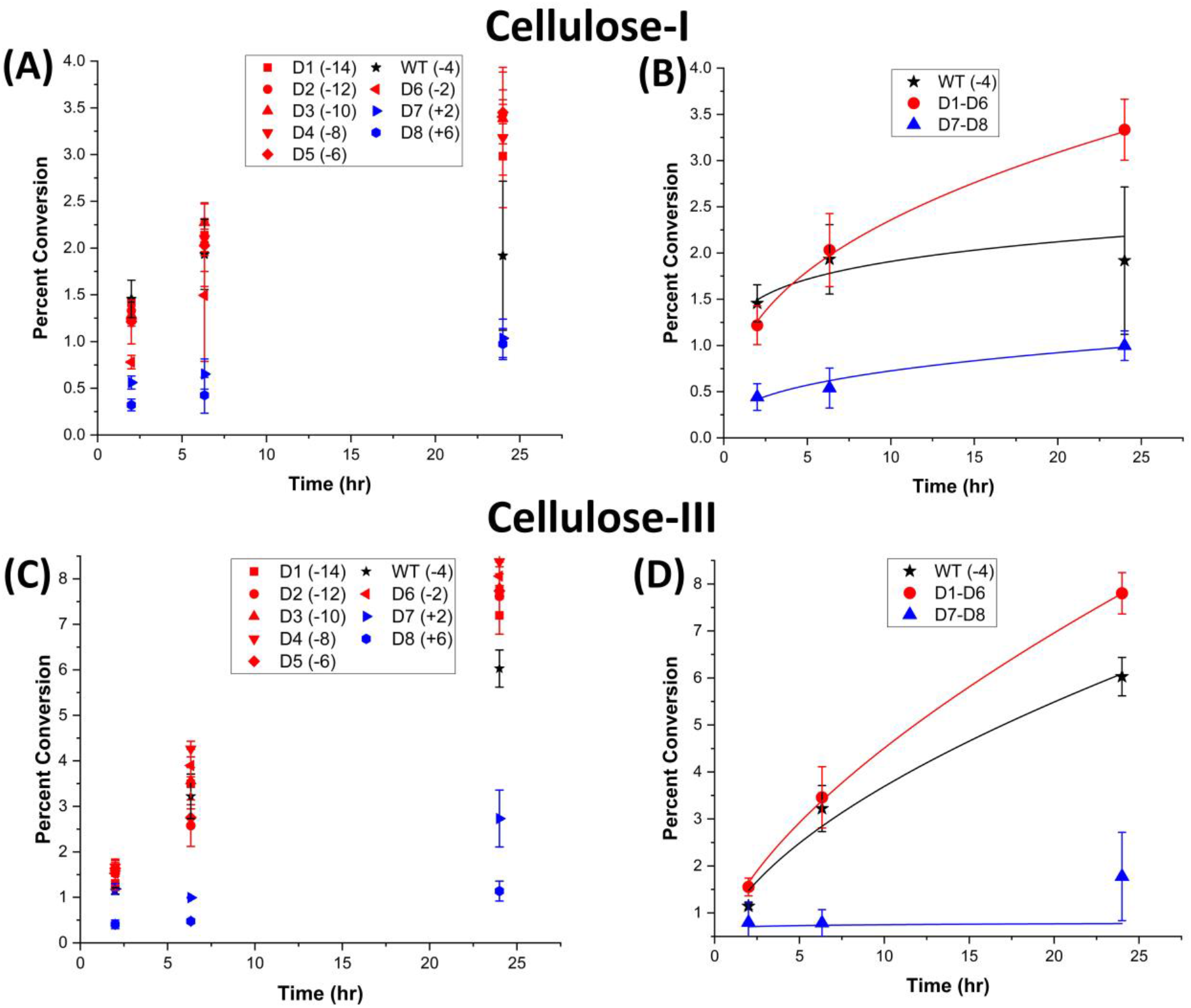
Negatively supercharged cellulases show improved hydrolysis yields on Avicel Cellulose-I and Cellulose-III. 40 μL of 100 g/L cellulose substrate was hydrolyzed using an enzyme loading of 120 nmol CBM2a Cel5A fusion enzyme per gram cellulose substrate supplemented with 12 nmol of β-glucosidase enzyme (10% of cellulase loading) per gram cellulose substrate for reaction times of 2hrs, 6hrs, and 24hrs. The solubilized reducing sugar concentrations in the supernatant after hydrolysis were determined by the DNS assay method. Percent conversion of biomass to glucose equivalents is reported on the Y-axis. (A) Percent conversion as a function of time (2 hr, 6 hr and 24 hr) for the hydrolysis of Avicel Cellulose-I by D1 – D8 CBM2a Cel5A and WT CBM2a Cel5A. Error bars represent standard deviation from the mean, based on at least 4 replicates. (B) Based on the data reported in (A), CBM2a Cel5A fusion constructs with negatively charged CBMs (D1 – D6) were grouped together and average hydrolysis yields were obtained for the group, with the error bars representing standard deviation from the mean. Similarly, CBM2a-Cel5A fusion constructs with positively charged CBMs (D7 and D8) were grouped together and average hydrolysis yields were obtained. Trend curves have been added to represent the kinetic profiles of the hydrolysis reaction. (C) Percent conversion as a function of time (2 hr, 6 hr and 24 hr) for the hydrolysis of Avicel Cellulose-III by D1 – D8 CBM2a Cel5A and WT CBM2a Cel5A. Error bars represent standard deviation from the mean, based on at least 4 replicates. (D) Based on the data reported in (C), CBM2a Cel5A fusion constructs with negatively charged CBMs (D1 – D6) were grouped together and average hydrolysis yields were obtained for the group, with the error bars representing standard deviation from the mean. Similarly, CBM2a-Cel5A fusion constructs with positively charged CBMs (D7 and D8) were grouped together and average hydrolysis yields were obtained. Trend curves have been added to represent the kinetic profiles of the hydrolysis reaction.

The two-parameter kinetic model fits for cellulose-I and cellulose-III are reported in **SI Appendix Table T6**. For cellulose-I, the negatively charged mutant enzyme group show ∼ 1.1 to 2.2-fold reduction in ‘A’ whereas b shows ∼ 1.9 to 4-fold improvement. Similar behavior is observed for positively charged group towards cellulose-I, although the reduction in A is more prominent (∼ 2.9-fold for D7 and ∼ 4-fold for D8) in this case. However, towards cellulose-III, the negatively charged mutants showed ∼ 1.8 to 2.3-fold improvement in A except for D2 and D5. This indicates a divergence in the behavior of mutants on cellulose-I versus cellulose-III although the overall trends in terms of improved hydrolysis yields are similar to that seen with pretreated biomass. For cellulose-I, the activity gains happen at a later stage potentially due to access to new substrate sites as indicated by the increase in b whereas for cellulose-III, the activity gains might arise from an improvement in the inherent activity of bound enzyme molecules as indicated by improvements seen in model parameter ‘A’.

Student’s T-tests were then performed between all pairs of mutants within a group (see **SI Appendix Table T7**), to understand individual differences. However, except for 5 pairs in the case of cellulose-I and 3 pairs in the case of cellulose-III (highlighted in red in **SI Appendix Table T7**), no statistically significant differences (p < 0.05) were observed. Hence, percent conversions for all the constructs in negatively charged group and positively charged group were averaged separately and represented in **Figure 3B** and **Figure 3D** for cellulose-I and cellulose-III, respectively.

In the case of cellulose-I (**Figure 3B**), the negatively charged sub-group (D1-D6) showed an improvement compared to the wild-type at all reaction time points with the maximum improvement (∼1.7-fold) over the wild-type enzyme being observed at 24 hours. A similar trend was observed for cellulose-III (**Figure 3D**) whereby the negatively charged sub-group (D1 – D6) showed a modest improvement (∼1.3-fold) over the wild-type at 24 hours. The modest improvement for cellulose-III compared to cellulose-I also partially explains the less significant impact of negative enzyme supercharging towards EA corn stover. However, the positively charged sub-group (D7 – D8) showed a 2 to 3.5-fold reduction in the case of cellulose-I and 1.5 to 4-fold reduction in the case of cellulose-III compared to the wild-type. To further understand how supercharged variants interact with cellulose substrates of different crystallinities, we also performed hydrolysis assays with phosphoric acid swollen cellulose (PASC) (reported in **SI Appendix Figure S3 – (A) for raw data and (B) for group means**). As seen in **SI Appendix Figure S3 (B)**, the negatively charged group did not show an improvement over the wild-type indicating that the enhanced activity trends seen for cellulose-I and cellulose-III might be unique to crystalline cellulosic substrates alone.

Overall, these results indicate an improvement in hydrolysis yields towards cellulose-I and cellulose-III, especially at longer reaction times such as 24 hours, albeit not at the same level of improvement in hydrolysis yields seen with pretreated biomass. Nevertheless, these results indicate that the improvement in specific towards crystalline cellulose could be one of the major factors contributing to the overall increase in biomass hydrolysis yields. Further, it was needed to be validated whether this improvement in activity towards cellulose arises from reduced CBM-induced non-productive binding. However, before proceeding to study binding, we wanted to test the hydrolysis yields towards crystalline cellulose in the presence of lignin.

### Positively supercharged cellulases show reduced crystalline cellulose hydrolysis yields in presence of exogenously added lignin

Lignin inhibition assays were performed by adding 1:2 (mass basis) of lignin and cellulose and incubating these synthetic substrates to mimic pretreated biomass along with enzymes for 24 hours. 24 hours was picked as the reaction time of choice, since the improvement in hydrolysis yields towards cellulose-I and cellulose-III for the negatively charged sub-group is evidenced at longer time points. As shown in **Figure 4**, both cellulose-I and cellulose-III showed similar hydrolysis yields for most enzyme designs except for D5 which showed greater activity towards cellulose-III. This assay resulted in larger error bars despite multiple trials possibly due to the presence of lignin in the slurry leading to lower signal-to-noise ratio during DNS reducing sugar assays. However, interestingly, there was a clear reduction in activity observed for the positively charged sub-group compared to the negatively charged sub-group, as highlighted in **Figure 4**.

**Figure 4:**
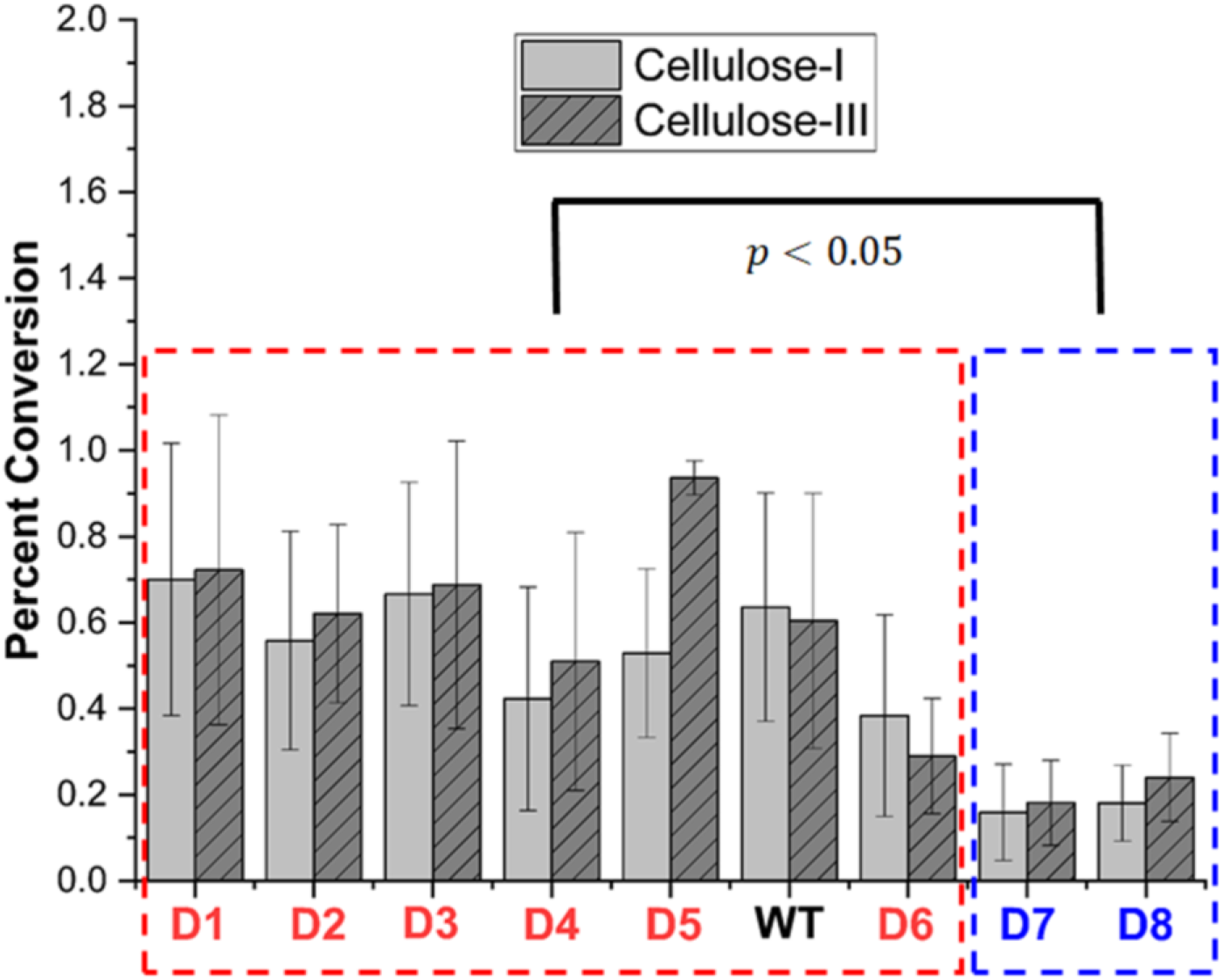
Negatively supercharged mutants perform better on Cellulose-I and Cellulose-III in the presence of exogenously added lignin. 20 μL of 100 g/L of Avicel Cellulose-I (light gray) or Avicel Cellulose-III (dark gray with lines) with addition of 80 μL of 25g/L dioxane extracted corn stover lignin was hydrolyzed using an enzyme loading of 120 nmol CBM2a Cel5A fusion enzyme per gram cellulose substrate and 12 nmol β-glucosidase per gram cellulose substrate for 24 hours. The solubilized reducing sugar concentrations in the supernatant after hydrolysis were determined by the DNS assay. Percent conversion of cellulosic substrate to glucose equivalents is reported on the Y-axis. T-test analysis (reported in the SI Appendix) was done to show that the activity of the fusion consutructs with negatively charged CBMs (red box) is statistically significantly different from that of the fusion constructs with positively charged CBMs.

Student’s T-tests were then conducted to validate these differences between a member of one group with every member of the other group (results summarized in **SI Appendix Table T8**). These results indicate that negatively charged CBM designs perform better in the presence of lignin. Overall, the results from cellulose hydrolysis assays and the lignin inhibition assays clearly indicate that the improved biomass hydrolysis yields reported in **Figure 2**, are a result of improved reactivity towards cellulose in the presence or absence of lignin. However, the negatively charged sub-group doesn’t show a clear improvement in hydrolysis yields compared to the wild-type in the presence of lignin, as might be expected from the results with pretreated biomass substrates discussed earlier. This trend could arise from a variety of factors including: 1. complex biomass ultrastructure cannot be reliably reproduced by a simple synthetic combination of cellulose and lignin, and 2. extraction of lignin leads to changes in its structure from that observed in pretreated biomass.

### QCM-D assays show reduced non-productive binding of negatively supercharged cellulases to both cellulose and lignin

Having validated the improved activity of negatively charged group of Cel5A-CBM2a constructs towards cellulose alone and also in the presence of lignin, we then studied the binding of CBM2a designs (attached to GFP) to crystalline cellulose and lignin using a QCM-D based method that we established previously^17,60^. Cellulose-I was picked as the crystalline cellulose substrate of choice for QCM-D binding analysis since it showed greater improvement in hydrolysis yield compared to cellulose-III. In addition, we have also established in a previous study^17^ that cellulose-I shows a greater level of off-pathway non-productive binding which can be alleviated by reducing CBM binding.

Raw QCM-D sensorgrams (as shown in **SI Appendix Figure S4** for cellulose and **SI Appendix Figure S5** for lignin) were fit to a mathematical model to obtain the maximum number of available binding sites and the desorption rate constant (*k*_*off*_) as discussed previously^17^. In the case of lignin, we computed the number of available binding sites and total percent protein recovered after binding, since lignin is known to mostly bind to proteins irreversibly as shown in previous studies^61^. As reported in Table 1, the negatively charged group (D1 – D6) shows a reduction in the number of binding sites (up to 1.5-fold) towards cellulose-I compared to the wild-type except for D5 and D6 towards cellulose-I. On the other hand, the positively charged CBM designs (D7 and D8) showed either comparable or slightly reduced number of binding sites as the wild-type. The desorption rate constant (*k*_*off*_) showed a slight increase for D1 – D4 (up to 1.3-fold as noticed for D2) and a reduction in the case of D5 and D6. As discussed in previous studies^17,64^, D1 – D4 show behavior representative of reduced non-productive binding to cellulose-I as indicated by a reduction in the number of binding sites and a slight increase in *k*_*off*_. On the other hand, the mutants in positively charged group (D7 and D8) show divergent behavior, with D7 showing an improvement in *k*_*off*_ whereas D8 shows a 10-fold reduction in *k*_*off*_.

**Table 1:** Binding of GFP-CBM2a fusion proteins to crystalline cellulose and lignin. Quartz crystal microbalance with dissipation (QCM-D) was used to study the binding of GFP-CBM2a designs to crystalline cellulose substrate (cellulose-I) and lignin. Number of binding sites (* 10^12^ molecules) and *k*_*off*_ (* 10^−3^ sec^-1^) were computed for cellulose-I whereas percent protein recovered was computed for lignin since binding was observed to be mostly irreversible. Experiments were performed in 2 replicates and the average along with standard deviation computed for these measurements which is reported here.

| CBM2a Design | Crystalline Cellulose |  | Lignin |  |
| --- | --- | --- | --- | --- |
| | Number of binding sites (* 10<sup>12</sup> molecules) | $k_{off}$ (* 10<sup>-3</sup> sec<sup>-1</sup>) | Number of binding sites (* 10<sup>12</sup> molecules) | Percent protein recovered |
| **D1 (-14)** | 95.00 ± 0.33 | 26.23 ± 0.27 | 23.46 ± 0.31 | 34.41 ± 2.84 |
| **D2 (-12)** | 151.65 ± 20.44 | 28.01 ± 1.83 | 25.42 ± 3.17 | 25.44 ± 0.27 |
| **D3 (-10)** | 115.19 ± 6.15 | 24.51 ± 5.46 | 18.83 ± 4.41 | 44.09 ± 16.16 |
| **D4 (-8)** | 122.77 ± 4.84 | 23.97 ± 1.27 | 26.11 ± 2.55 | 32.35 ± 5.54 |
| **D5 (-6)** | 210.54 ± 5.23 | 5.95 ± 0.78 | 41.05 ± 4.24 | 38.70 ± 1.41 |
| **WT (-4)** | 143.72 ± 9.11 | 21.06 ± 0.91 | 33.18 ± 1.02 | 19.55 ± 1.92 |
| **D6 (-2)** | 223.61 ± 11.96 | 18.63 ± 0.93 | 30.44 ± 2.18 | 49.22 ± 3.25 |
| **D7 (+2)** | 128.68 ± 5.41 | 26.06 ± 0.16 | 35.63 ± 2.43 | 31.52 ± 9.10 |
| **D8 (+6)** | 137.69 ± 5.38 | 2.02 ± 0.04 | 37.24 ± 1.75 | 32.18 ± 0.00 |

In the case of lignin, all mutants in the negatively charged group (D1 – D6) show a reduction (up to 1.7-fold) in the number of binding sites compared to the wild-type except for D5. However, the positively charged mutants (D7 and D8) show an increase in the number of binding sites. This trend is in accordance with the original hypothesis that negative supercharging could lead to a reduction in binding to lignin due to electrostatic repulsions as outlined in previous studies from our group as well^28,29^. We also characterized the percentage of protein recovered during unbinding from lignin and surprisingly all mutants showed an improvement in protein recovery when compared to the wild-type. A reduction in number of binding sites and an improvement in protein recovery is highly desirable because irreversible attachment of enzymes to lignin via CBM would lead to enzyme deactivation.

Overall, relating the results from QCM-D binding assays with hydrolysis assays on pretreated biomass and cellulose, it is likely that the improved hydrolytic activity of D1 – D4 CBM2a Cel5A towards pretreated biomass arises from reduced non-productive binding to cellulose, caused by a reduction in the number of binding sites and an increase in *k*_*off*_ and also likely from reduced non-productive binding to lignin. This definition of non-productive binding to cellulose is in accordance with the explanation from previous studies as well^64^. However, D5 and D6 CBM2a Cel5A show trends divergent from the rest of the negatively charged group in regards to binding to cellulose, with an increase in the number of binding sites coupled with a decrease in the desorption rate constant (*k*_*off*_). It is likely that the new substrate binding sites made accessible by the mutations on D5 and D6 CBM2a Cel5A are more prone to hydrolysis but validation of this hypothesis requires further structural and biophysical investigation. On the other hand, although the lignin inhibition assay results for D7 and D8 CBM2a Cel5A can be rationalized based on the increased binding to lignin, we still cannot explain the reduced hydrolytic activity towards cellulose-I and cellulose-III and thereby pretreated biomass. To further address these unanswered questions, we chose to measure the thermal stability of all mutants since it has a significant impact on biomass hydrolysis at higher temperatures.

### Supercharged cellulase designs retain thermal stability

We performed thermal shift assays on all CBM2a-Cel5A fusion constructs using a SYPRO orange-based procedure established previously^29^. The raw data (reported in **SI Appendix Figure S6**) was analyzed using an open-source tool called SimpleDSFViewer^58^, to obtain melting temperatures (reported in **Table 2**). All mutant designs exhibit melting temperatures similar to the wild-type enzyme with minor fluctuations within a narrow range of 1 °C. However, these results indicate the melting temperature for the entire CBM2a-Cel5A enzyme whereas the variation due to supercharging might only be confined to the CBM portion of the enzyme. Hence, we are currently exploring methods to obtain the melting temperatures of the CBM alone, to provide further mechanistic credence to the binding and hydrolysis results reported here.

**Table 2.** Thermal shift assays show negligible impact of mutations on CBM2a-Cel5A cellulase melting temperatures. Thermal shift assays were conducted using 5 µl 200X SYPRO reagent, 5 µl 0.5 M sodium acetate buffer (pH 5.5), 5 µM enzyme dilution and deionized water to make up to 50 µl. The mixtures were exposed to a temperature ramp from 25 °C to 99 °C at a rate of 0.04 °C per second using QuantStudio 3 real-time PCR equipment and fluorescence measured (excitation – 470 nm; emission – 520 nm). Melting temperatures (T_m_) were obtained from each individual trace and averages with standard deviations reported here.

| Design | Net charge | T<sub>m</sub> (°C) |
| --- | --- | --- |
| D1 | -14 | 73.74 ± 0.08 |
| D2 | -12 | 73.36 ± 0.11 |
| D3 | -10 | 73.62 ± 0.15 |
| D4 | -8 | 74.84 ± 0.03 |
| D5 | -6 | 74.32 ± 0.12 |
| WT | -4 | 74.02 ± 0.01 |
| D6 | -2 | 73.54 ± 0.17 |
| D7 | +2 | 73.67 ± 0.07 |
| D8 | +6 | 74.22 ± 0.11 |

## CONCLUSIONS

Carbohydrate-binding modules (CBMs) play a crucial role in targeting appended glycoside hydrolase enzymes to plant cell wall polymers such as cellulose and hemicellulose^65–68^. However, recent studies have shown that CBMs can also play a role in non-productive binding of appended cellulase catalytic domains to cellulose surface^17,69,70^. In addition, CBMs can bind non-productively to lignin via hydrophobic interactions, leading to deactivation of the enzymes on biomass surface^42,61^. To address these bottlenecks, a previous study from our lab has used selective supercharging of cellulase enzymes to reduce lignin inhibition although the mechanistic details were yet to be elucidated^29^. In this study, we expanded the supercharging approach to address mechanistic questions surrounding the impact of CBM supercharging on the hydrolysis of real-world pretreated biomass substrates.

In summary, we established that the improvement in hydrolytic activity (up to 3.3-fold) observed by negatively supercharging CBMs could arise from a combination of factors including reduced non-productive binding to cellulose leading to improvement in cellulolytic activity (up to 1.7-fold), reduced non-productive binding to lignin (up to 1.8-fold), increased reversible binding to lignin (up to 2.2-fold), while retaining the thermal stability of these fusion enzymes. It is likely that the reduction in non-productive binding to cellulose and lignin upon negative supercharging, arises from the negative zeta potential of cellulose and lignin leading to electrostatic repulsions between their surfaces and the negatively charged protein surface, as previously hypothesized^29^. Given the promising nature of this approach to improve cellulase enzyme performance, it remains to be seen how supercharging impacts various families of glycoside hydrolase enzymes secreted by biomass-degrading microbes. In addition, future work should address the application of supercharging to selectively fine-tune inter-enzyme interactions to enable increased enzyme synergism, inspired by recent work using supercharging to enable self-assembly of electrostatically complementary proteins^35^.

## Supporting information

SI Appendix Sequences

## ACKNOWLEDGEMENTS

The authors acknowledge support from the NSF CBET (Award 1604421 and Career Award 1846797), ORAU Ralph E. Powe Award, Rutgers Global Grant, Rutgers Division of Continuing Studies, Rutgers School of Engineering, and the Great Lakes Bioenergy Research Center (DOE BER Office of Science DE-FC02-07ER64494). The authors thank Nathan Kruer-Zerhusen (from late Prof. David Wilson’s lab) for kindly providing the *Thermobifida fusca* strain and original Cel5A plasmid DNA. Special thanks to Sarvada Chipkar and Dr. Rebecca Ong at Michigan Technological University, for conducting ammonia pretreatment to provide us with cellulose-III and AFEX corn stover. SPSC would also like to thank Dr. Leonardo Da Sousa (Dale lab at Michigan State University) for conducting ammonia pretreatment to generate cellulose III and EA corn stover that was used here in this study as well. BN would like to thank Patrick Doran (Boston University) for his help initially with setting up protocols for computational protein design using Rosetta. JMY acknowledges support from the Department of Energy, Office of Energy Efficiency and Renewable Energy (EERE) under agreement no. 28598.

## SUPPLEMENTARY SECTION

See supplementary information pdf titled ‘SI_Appendix.pdf’ for supporting results relevant to this study. See word document titled ‘SI_Appendix_Sequences.docx’, which has the full nucleotide sequences for pEC-CBM2a-Cel5A and pEC-GFP-CBM2a and highlights the mutation sites in CBM2a amino acid sequence.

## Graphical abstract

Negatively supercharged cellulases show improved biomass hydrolysis activity due to reduced non-productive binding to lignin and crystalline cellulose

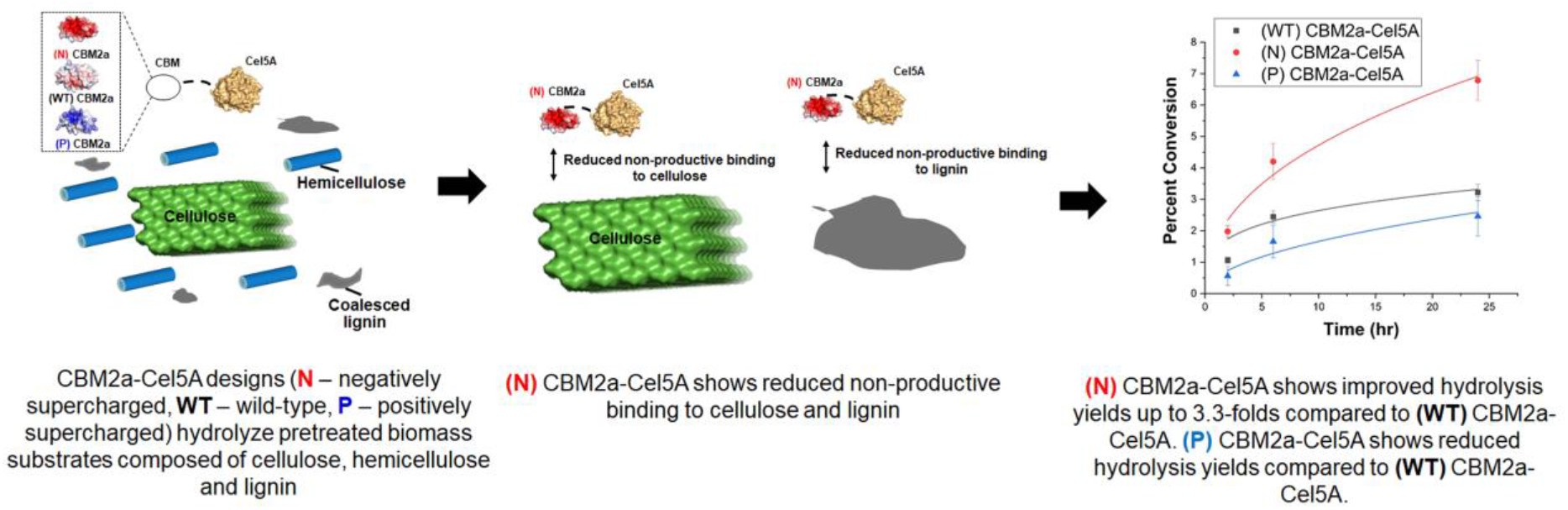

