## Supplementary material for "Supercharged Cellulases Show Reduced Non-Productive Binding, But Enhanced Activity, on Pretreated Lignocellulosic Biomass": SI Appendix

**Number of SI pages in current PDF (SI-Supplementary Information): 17**

**Number of SI figures in current PDF (SI-Supplementary Information): 6**

**Number of SI tables in current PDF (SI-Supplementary Information): 8**

**Number of SI files (including current PDF): 2**

[#] Equal contribution

| **CBM2a Construct** | **Nucleotide Sequence** |
|---|---|
| WT | GCCGGCTGCTCGGTGGACTACACGGTCAACTCCTGGGGTACCGGGTTCACC<br>GCCAACGTCACCATCACCAACCTCGGCAGTGCGATCAACGGCTGGACCCTG<br>GAGTGGGACTTCCCCGGCAACCAGCAGGTGACCAACCTGTGGAACGGGACC<br>TACACCCAGTCCGGGCAGCACGTGTCGGTCAGCAACGCCCCGTACAACGCC<br>TCCATCCCGGCCAACGGAACGGTTGAGTTCGGGTTCAACGGCTCCTACTCGG<br>GCAGCAACGACATCCCCTCCTCCTTCAAGCTGAACGGGGTTACCTGC |
| D1 | GCCGGTTGCTCTGTTGATTATACCGTCAATTCTTGGGGTACGGGTTTTACAG<br>CGAACGTAACCATTACTAACCTTGGAGACGCCATCGAGGGGTGGGATTTAG<br>AGTGGGACTTTCCAGGTGACCAGCAGGTAACTAACCTTTGGAACGGAGAAT<br>ACGAGCAAGACGGTGAACACGTCAGTGTGGAGAATGCGCCTTACAATGCTT<br>CGATCCCAGCTAATGGAACCGTCGAGTTCGGGTTTAACGGTGAATACAGTG<br>GCTCCAACGACATCCCTAGTTCGTTCAAGTTGAATGGAGTGACCTGC |
| D2 | GCCGGCTGCTCGGTGGACTACACGGTCAACTCCTGGGGTACCGGGTTCACT<br>GCGAATGTTACCATTACAAACTTGGGAGATGCCATCAACGGATGGGATCTT<br>GAATGGGACTTCCCCGGTGACCAACAAGTTACCAATCTGTGGAACGGAGAA<br>TATGAGCAGAGTGGCGAGCATGTAAGCGTTGAAAATGCTCCTTACAATGCT<br>TCGATTCCTGCTAACGGTACGGTGGAGTTCGGTTTTAATGGAGAGTACTCTG<br>GCAGCAACGACATCCCCTCCTCCTTCAAGCTGAACGGGGTTACCTGC |
| D3 | GCCGGCTGCTCGGTGGACTACACGGTCAACTCCTGGGGTACCGGGTTCACG<br>GCCAATGTAACCATTACAAATTTGGGTGACGCCATCAACGGTTGGGACCTT<br>GAATGGGATTTTCCCGGAGATCAACAGGTAACAAACCTGTGGAATGGTGAA<br>TATGAACAGTCCGGGGAGCATGTGTCGGTGAGTAACGCTCCCTATAATGCTT<br>CCATCCCGGCGAATGGGACTGTTGAGTTTGGGTTTAACGGAAGTTATAGCG<br>GCAGCAACGACATCCCCTCCTCCTTCAAGCTGAACGGGGTTACCTGC |
| D4 | GCCGGCTGCTCGGTGGACTACACGGTCAACTCCTGGGGTACCGGGTTCACT<br>GCGAACGTGACGATCACAAATTTAGGCTCGGCTATTAACGGGTGGGATTTG<br>GAATGGGATTTTCCTGGAGATCAACAGGTTACCAACTTATGGAACGGTGAG<br>TACGAACAGTCAGGTCAACACGTAAGCGTATCTAATGCACCTTACAATGCC<br>TCAATTCCAGCCAATGGGACTGTAGAATTTGGGTTTAATGGCAGTTATTCTG<br>GCAGCAACGACATCCCCTCCTCCTTCAAGCTGAACGGGGTTACCTGC |
| D5 | GCCGGCTGCTCGGTGGACTACACGGTCAACTCCTGGGGTACCGGGTTCACA<br>GCAAATGTAACAATCACTAATTTAGGCTCCGCTATTAACGGGTGGACCTTGG<br>AATGGGATTTCCCCGGTGACCAGCAGGTAACGAACCTTTGGAACGGTGAAT<br>ACACGCAGTCCGGGCAACACGTAAGTGTCTCCAATGCCCCTTACAATGCAT<br>CTATCCCTGCTAATGGAACCGTCGAGTTCGGCTTTAACGGTTCATACTCCGG<br>CAGCAACGACATCCCCTCCTCCTTCAAGCTGAACGGGGTTACCTGC |
| D6 | GCCGGCTGCTCGGTGGACTACACGGTCAACTCCTGGGGTACCGGGTTCACA<br>GCGAACGTAACAATCACAAATTTAGGATCTGCAATCAATGGCTGGAAGCTG<br>GAGTGGGATTTTCCGGGAAATCAACAAGTGACCAACTTGTGGAACGGCACC<br>TATAAGCAATCCGGCCAGCACGTATCAGTGTCGAACGCCCCCTATAATGCTT<br>CGATCCCGGCCAATGGTACGGTGGAGTTCGGATTTAACGGGTCATACTCTG<br>GCAGCAACGACATCCCCTCCTCCTTCAAGCTGAACGGGGTTACCTGC |
| D7 | GCCGGCTGCTCGGTGGACTACACGGTCAACTCCTGGGGTACCGGGTTCACT<br>GCAAACGTAACCATCACCAACTTGGGCCGCGCTATCAATGGTTGGAAGCTT<br>GAGTGGGATTTCCCCGGCAACCAGCAGGTAACAAATCTTTGGAACGGGACC<br>TATAAGCAGTCAGGGAAACACGTAAGCGTGAAAAATGCGCCTTACAACGCA<br>TCAATCCCAGCTAATGGCACAGTAGAGTTTGGATTTAACGGTAAATACTCCG<br>GCAGCAACGACATCCCCTCCTCCTTCAAGCTGAACGGGGTTACCTGC |

| | |
|---|---|
| D8 | GCCGGGTGTAGTGTGGATTACACTGTGAATTCCTGGGGTACAGGATTCACG<br>GCCAATGTAACGATAACAAACTTAGGACGGGCTATTAAAGGTTGGAAGCTG<br>GAGTGGGACTTCCCTGGTAAGCAACAAGTCACGAATCTGTGGAATGGACGT<br>TATAAGCAAAAAGGGAAGCATGTTTCGGTCAAAAATGCACCGTATAACGCT<br>AGTATCCCTGCAAACGGGACAGTGGAATTCGGTTTTAATGGCAAGTATAGC<br>GGGTCGAATGATATACCATCTTCATTTAAGTTAAATGGAGTCACTTGC |

**Table T1: Nucleotide sequences of supercharged CBM2a designs including the wild-type (labeled WT).** The nucleotide sequence of each CBM2a design reported here, can be swapped with the CBM2a wild-type reported in Table T1 to obtain the respective pEC-CBM2a-Cel5A and pEC-GFP-CBM2a versions.

| | Mutation to **K** | Mutation to **R** | Mutation to **D** | Mutation to **E** | Average mutation energy score (K, R, D, E) |
|---|---|---|---|---|---|
| **Cluster 1** | | | | | |
| N39 | 0.972 | 1.339 | -1.545 | -0.372 | 0.0985 |
| T51 | -0.319 | -1.632 | 0.507 | -0.391 | -0.45875 |
| Q55 | -0.964 | -0.648 | -0.185 | -0.308 | -0.52625 |
| S81 | -0.804 | -0.173 | 0.159 | -1.979 | -0.69925 |
| S53 | -1.236 | -0.886 | -1.314 | -0.818 | -1.0635 |
| S83 | -0.263 | 1.587 | -0.859 | -0.386 | 0.01975 |
| **Cluster 2** | | | | | |
| T49 | -0.925 | -1.243 | 0.01 | -0.628 | -0.6965 |
| T31 | -1.96 | -1.625 | 0.246 | 0.362 | -0.74425 |
| S25 | -0.503 | -1.424 | -1.112 | -0.31 | -0.83725 |
| S60 | -0.925 | -0.707 | -1.133 | -1.008 | -0.94325 |
| N28 | -1.944 | -1.806 | -0.413 | -2.676 | -1.70975 |

**Table T2**: **Rosetta mutation energy scores for polar uncharged surface residues based after multiple fast relax operations until the energy score equilibrates.** K, R, D, E mutations were scored for each polar uncharged residue using Rosetta and an average mutation energy score obtained. Based on the homology model of wild-type CBM2a, these residues were organized into two spatially distinct clusters.

| Design | Net Charge | Mutations |
|---|---|---|
| D1 | -14 | N39D, T49E, T51E, T31D, Q55E, S25D, S81E, S60E, S53D, N28E |
| D2 | -12 | N39D, T49E, T51E, T31D, Q55E, S25D, S81E, S60E |
| D3 | -10 | N39D, T49E, T51E, T31D, Q55E, S25D |
| D4 | -8 | N39D, T49E, T51E, T31D |
| D5 | -6 | N39D, T49E |
| D6 | -2 | T51K, T31K |
| D7 | +2 | T51K, T31K, Q55K, S25R, S81K, S60K |
| D8 | +6 | T51K, T31K, Q55K, S25R, S81K, S60K, S53K, N28K, N39K, T49R |

**Table T3: Mutations on wild-type CBM2a to generate supercharged mutant designs.** Based on the average energy scores computed in previous table T1, surface residues were either mutated to a negatively charged residue (D, E) or a positively charged residue (R, K) to obtain a target net charge. This table reports the list of mutations necessary to generate a certain CBM2a mutant design (D1-D8) from CBM2a wild-type.

| | **AFEX Corn Stover** | | | **EA Corn Stover** | | |
|---|---|---|---|---|---|---|
| **Enzyme Design** | **A** | **b** | **Adjusted R-Square** | **A** | **b** | **Adjusted R-Square** |
| **D1 (-14)** | 3.20 ± 1.06 | 0.32 ± 0.09 | 0.99 | 3.53 ± 0.97 | 0.42 ± 0.07 | 0.99 |
| **D2 (-12)** | 2.09 ± 0.23 | 0.42 ± 0.04 | 0.99 | 2.27 ± 0.49 | 0.51 ± 0.06 | 0.99 |
| **D3 (-10)** | 2.01 ± 0.14 | 0.39 ± 0.02 | 0.99 | 2.79 ± 0.26 | 0.44 ± 0.03 | 0.99 |
| **D4 (-8)** | 2.30 ± 0.22 | 0.37 ± 0.03 | 0.99 | 2.86 ± 0.15 | 0.47 ± 0.01 | 0.99 |
| **D5 (-6)** | 2.40 ± 0.47 | 0.35 ± 0.06 | 0.99 | 2.83 ± 0.21 | 0.46 ± 0.02 | 0.99 |
| **WT (-4)** | 0.97 ± 0.19 | 0.33 ± 0.06 | 0.99 | 1.54 ± 0.54 | 0.44 ± 0.10 | 0.99 |
| **D6 (-2)** | 2.02 ± 0.23 | 0.36 ± 0.03 | 0.99 | 3.53 ± 0.60 | 0.36 ± 0.05 | 0.99 |
| **D7 (+2)** | 0.68 ± 0.04 | 0.34 ± 0.02 | 0.99 | 1.69 ± 0.35 | 1.69 ± 0.36 | 0.99 |
| **D8 (+6)** | 0.21 ± 0.13 | 0.55 ± 0.16 | 0.99 | 0.47 ± 0.04 | 0.47 ± 0.15 | 0.99 |
| **D1-D6** | 2.19 ± 0.40 | 0.37 ± 0.06 | 0.99 | 2.76 ± 0.41 | 2.77 ± 0.41 | 0.99 |
| **D7-D8** | 0.51 ± 0.11 | 0.39 ± 0.05 | 0.99 | 0.89 ± 0.24 | 0.89 ± 0.24 | 0.99 |

**Table T4: Kinetic parameter fits for biomass hydrolysis by CBM2a-Cel5A fusion constructs.** Reaction progress curves for hydrolysis of AFEX and EA pretreated corn stover (reported in **Fig. 2** of main manuscript) by CBM2a-Cel5A fusion constructs was fit to a power curve represented by the formula $Percent\ conversion = A * time^b$. Curve fitting was done in Origin and the averages, standard errors from mean for each parameter are reported here. WT refers to wild-type.

| | | AFEX Corn Stover | | | EA Corn Stover | | |
|---|---|---|---|---|---|---|---|
| **Construct 1** | **Construct 2** | **2 hrs** | **6 hrs** | **24 hrs** | **2 hrs** | **6 hrs** | **24 hrs** |
| D1 | D2 | 0.232 | 0.044 | 0.062 | 0.728 | 0.001 | 0.040 |
| D1 | D3 | 0.210 | 0.021 | 0.015 | 0.839 | 0.002 | 0.036 |
| D1 | D4 | 0.736 | 0.053 | 0.007 | 0.645 | 0.001 | 0.356 |
| D1 | D5 | 0.152 | 0.055 | 0.546 | 0.154 | 0.005 | 0.710 |
| D1 | D6 | 0.013 | 0.017 | 0.136 | 0.426 | 0.002 | 0.010 |
| D2 | D3 | 0.776 | 0.093 | 0.684 | 0.337 | 0.129 | 0.713 |
| D2 | D4 | 0.201 | 0.512 | 0.907 | 0.827 | 0.765 | 0.015 |
| D2 | D5 | 0.041 | 0.343 | 0.110 | 0.050 | 0.734 | 0.029 |
| D2 | D6 | 0.145 | 0.342 | 0.322 | 0.066 | 0.182 | 0.153 |
| D3 | D4 | 0.222 | 0.060 | 0.657 | 0.386 | 0.406 | 0.012 |
| D3 | D5 | 0.011 | 0.045 | 0.086 | 0.008 | 0.215 | 0.038 |
| D3 | D6 | 0.025 | 0.891 | 0.381 | 0.216 | 0.010 | 0.300 |
| D4 | D5 | 0.398 | 0.607 | 0.054 | 0.214 | 0.597 | 0.172 |
| D4 | D6 | 0.044 | 0.271 | 0.250 | 0.159 | 0.209 | 0.004 |
| D5 | D6 | 0.001 | 0.237 | 0.355 | 0.000 | 0.569 | 0.015 |
| D7 | D8 | 0.004 | 0.026 | 0.834 | 0.003 | 0.000 | 0.012 |

**Table T5: p-values from Student's T-test for comparison of biomass hydrolysis yields from CBM2a-Cel5A fusion constructs.** Student's T-test was conducted to compare biomass hydrolysis yields of every pair in the negatively charged group (D1 – D6) and every pair in the positively charged group (D7 – D8). The raw data is reported in **Fig. 2A** and **2C** of the main manuscript.

| | Cellulose-I | | | Cellulose-III | | |
|---|---|---|---|---|---|---|
| **Enzyme** | **A** | **b** | **Adjusted R-Square** | **A** | **b** | **Adjusted R-Square** |
| **D1 (-14)** | 1.23 ± 0.10 | 0.28 ± 0.03 | 0.99 | 1.10 ± 0.25 | 0.59 ± 0.08 | 0.99 |
| **D2 (-12)** | 1.02 ± 0.01 | 0.38 ± 0.004 | 0.99 | 0.76 ± 0.20 | 0.73 ± 0.08 | 0.99 |
| **D3 (-10)** | 1.13 ± 0.17 | 0.35 ± 0.05 | 0.99 | 1.16 ± 0.07 | 0.60 ± 0.02 | 0.99 |
| **D4 (-8)** | 1.02 ± 0.14 | 0.38 ± 0.07 | 0.99 | 1.53 ± 0.28 | 0.54 ± 0.06 | 0.99 |
| **D5 (-6)** | 0.93 ± 0.03 | 0.42 ± 0.01 | 0.99 | 0.80 ± 0.18 | 0.71 ± 0.07 | 0.99 |
| **WT (-4)** | 1.34 ± 0.16 | 0.15 ± 0.08 | 0.99 | 0.99 ± 0.34 | 0.57 ± 0.11 | 0.99 |
| **D6 (-2)** | 0.51 ± 0.01 | 0.60 ± 0.01 | 0.99 | 1.27 ± 0.23 | 0.58 ± 0.06 | 0.99 |
| **D7 (+2)** | 0.46 ± 0.05 | 0.24 ± 0.05 | 0.99 | 0.99 ± 0.61 | 0 ± 0.33 | 0.96 |
| **D8 (+6)** | 0.22 ± 0.04 | 0.46 ± 0.07 | 0.99 | 0.18 ± 0.08 | 0.56 ± 0.21 | 0.99 |
| **D1-D6** | 0.96 ± 0.05 | 0.39 ± 0.02 | 0.99 | 1.07 ± 0.05 | 0.63 ± 0.01 | 0.99 |
| **D7-D8** | 0.33 ± 0.06 | 0.35 ± 0.07 | 0.99 | 0.69 ± 0.46 | 0.04 ± 0.37 | 0.95 |

**Table T6: Kinetic parameter fits for crystalline cellulose hydrolysis by CBM2a-Cel5A fusion constructs.** Reaction progress curves for hydrolysis of cellulose-I and cellulose-III (reported in **Fig. 3** of main manuscript) by CBM2a-Cel5A fusion constructs was fit to a power curve represented by the formula $Percent\ conversion = A * time^b$. Curve fitting was done in Origin and the averages, standard errors from mean for each parameter are reported here. WT refers to wild-type.

| | | Cellulose-I | | | Cellulose-III | | |
|---|---|---|---|---|---|---|---|
| **Construct 1** | **Construct 2** | **2 hrs** | **6 hrs** | **24 hrs** | **2 hrs** | **6 hrs** | **24 hrs** |
| D1 | D2 | 0.158 | 0.716 | 0.152 | 0.385 | 0.162 | 0.382 |
| D1 | D3 | 0.881 | 0.129 | 0.603 | 0.213 | 0.173 | 0.449 |
| D1 | D4 | 0.264 | 0.697 | 0.770 | 0.147 | 0.158 | 0.094 |
| D1 | D5 | 0.014 | 0.610 | 0.201 | 0.150 | 0.157 | 0.306 |
| D1 | D6 | 0.049 | 0.410 | 0.152 | 0.448 | 0.330 | 0.167 |
| D2 | D3 | 0.507 | 0.437 | 0.729 | 0.049 | 0.045 | 0.065 |
| D2 | D4 | 0.467 | 0.813 | 0.719 | 0.560 | 0.090 | 0.143 |
| D2 | D5 | 0.074 | 0.975 | 0.947 | 0.625 | 0.690 | 0.258 |
| D2 | D6 | 0.009 | 0.452 | 0.877 | 1.000 | 0.085 | 0.055 |
| D3 | D4 | 0.653 | 0.171 | 0.770 | 0.854 | 0.062 | 0.226 |
| D3 | D5 | 0.319 | 0.344 | 0.819 | 0.939 | 0.066 | 0.874 |
| D3 | D6 | 0.008 | 0.353 | 0.928 | 0.665 | 0.240 | 0.514 |
| D4 | D5 | 0.983 | 0.721 | 0.705 | 0.828 | 0.015 | 0.153 |
| D4 | D6 | 0.074 | 0.428 | 0.751 | 0.611 | 0.230 | 0.299 |
| D5 | D6 | 0.033 | 0.463 | 0.879 | 0.678 | 0.019 | 0.099 |
| D7 | D8 | 0.010 | 0.196 | 0.760 | 0.001 | 0.001 | 0.150 |

**Table T7: p-values from Student's T-test for comparison of crystalline cellulose hydrolysis yields from CBM2a-Cel5A fusion constructs.** Student's T-test was conducted to compare biomass hydrolysis yields of every pair in the negatively charged group (D1 – D6) and every pair in the positively charged group (D7 – D8). The raw data is reported in **Fig. 3A** and **3C** of the main manuscript.

| | Cellulose-I | | Cellulose-III | |
|---|---|---|---|---|
| | **Comparison to D7** | **Comparison to D8** | **Comparison to D7** | **Comparison to D8** |
| **D1** | 0.001 | 0.002 | 0.003 | 0.006 |
| **D2** | 0.002 | 0.003 | 0.000 | 0.001 |
| **D3** | 0.000 | 0.001 | 0.003 | 0.006 |
| **D4** | 0.022 | 0.031 | 0.017 | 0.040 |
| **D5** | 0.000 | 0.001 | 0.000 | 0.000 |
| **D6** | 0.028 | 0.040 | 0.083 | 0.413 |
| **WT** | 0.001 | 0.001 | 0.004 | 0.010 |

**Table T8**: **p-values from Student's T-test for comparison of hydrolysis yields in the presence of lignin.** Student's T-test was conducted to compare the hydrolysis yields of all the constructs in negatively charged group (D1 – D6) to all the constructs in positively charged group (D7 – D8). These results reveal statistically significant differences between any pair of constructs where one mutant construct is derived from the positively charged group

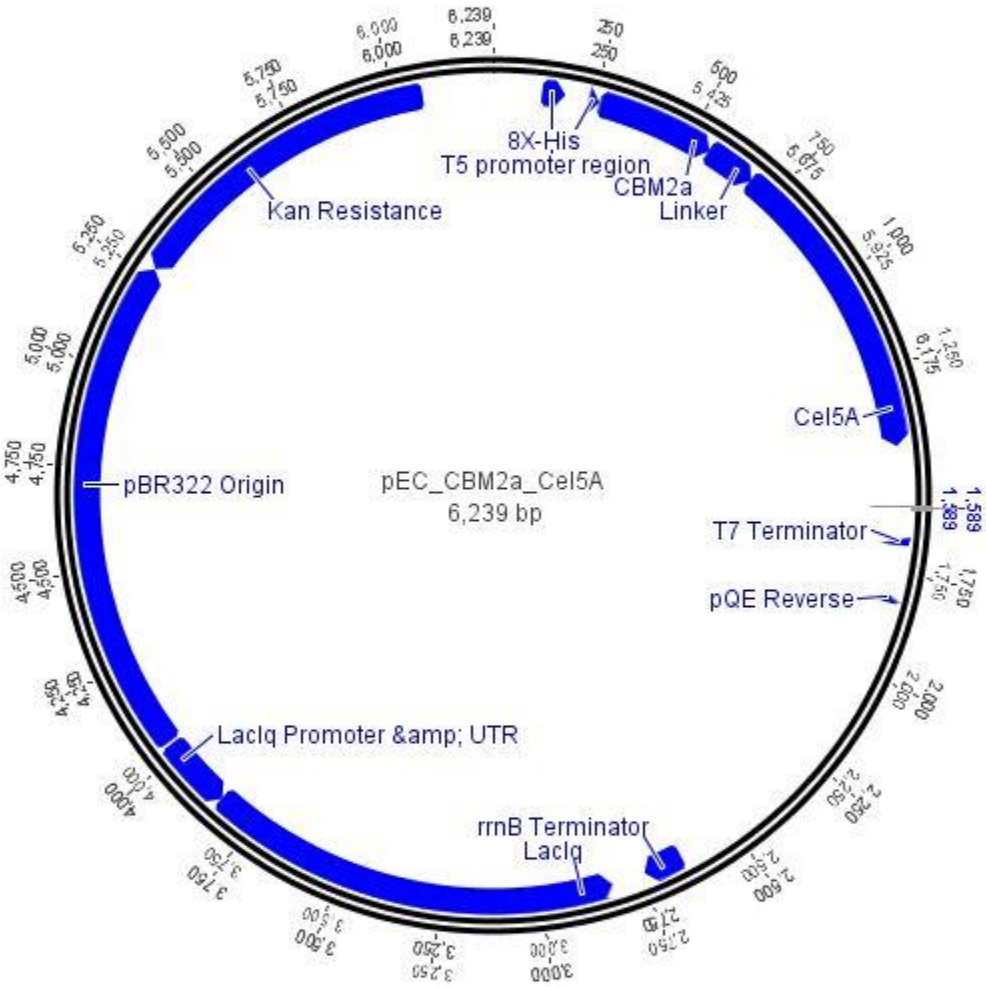


**Figure S1**: **Plasmid map for pEC-CBM2a-Cel5A** Open reading frame consisting of 8X Histidine tag (labeled 8X-His) followed by CBM2a wild-type, linker and Cel5A is highlighted. pEC plasmid backbone consists of T5 promoter region and T7 terminator region as described elsewhere[1]. Other elements highlighted here are: pBR322 origin of replication, LacI promoter and gene to enable lactose-inducible expression.

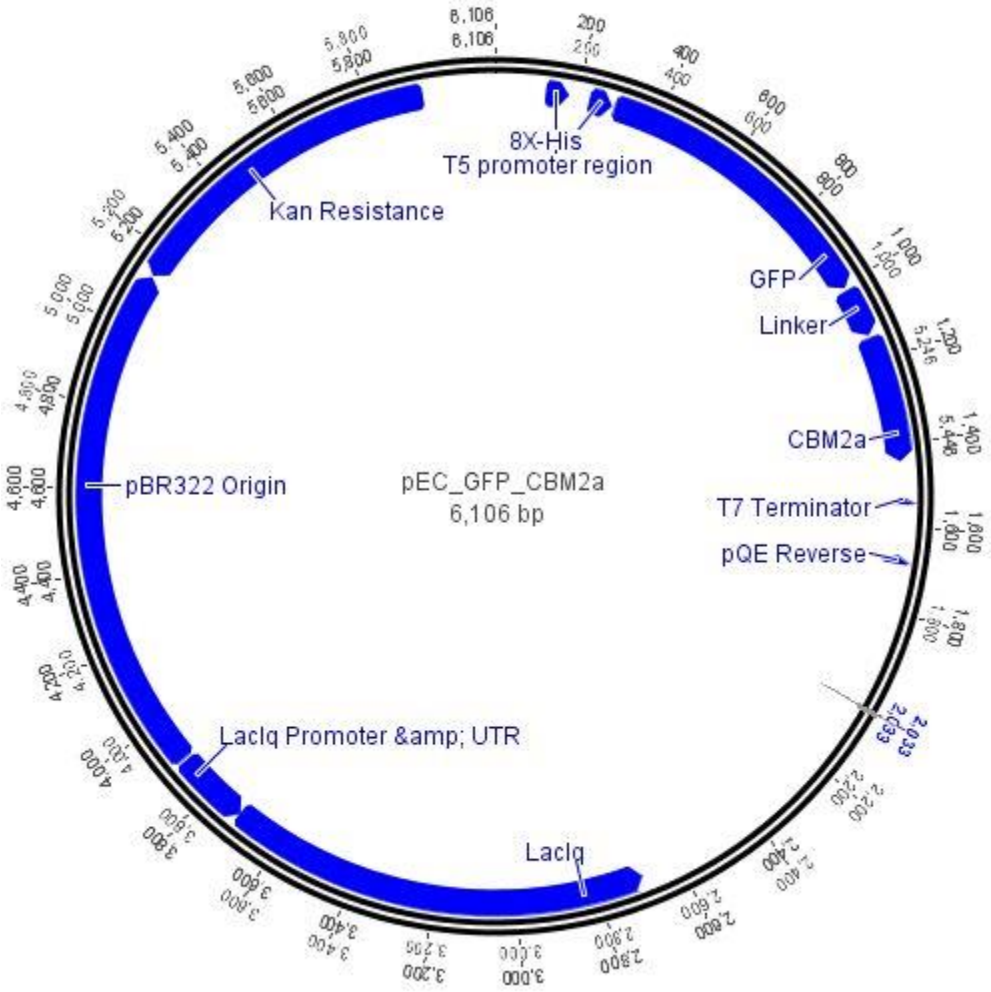


**Figure S2**: **Plasmid map for pEC-GFP-CBM2a:** Open reading frame consisting of 8X Histidine tag (labeled 8X-His) followed by GFP, linker and CBM2a wild-type is highlighted. pEC plasmid backbone consists of T5 promoter region and T7 terminator region as described elsewhere[1]. Other elements highlighted here are: pBR322 origin of replication, LacI promoter and gene to enable lactose-inducible expression.

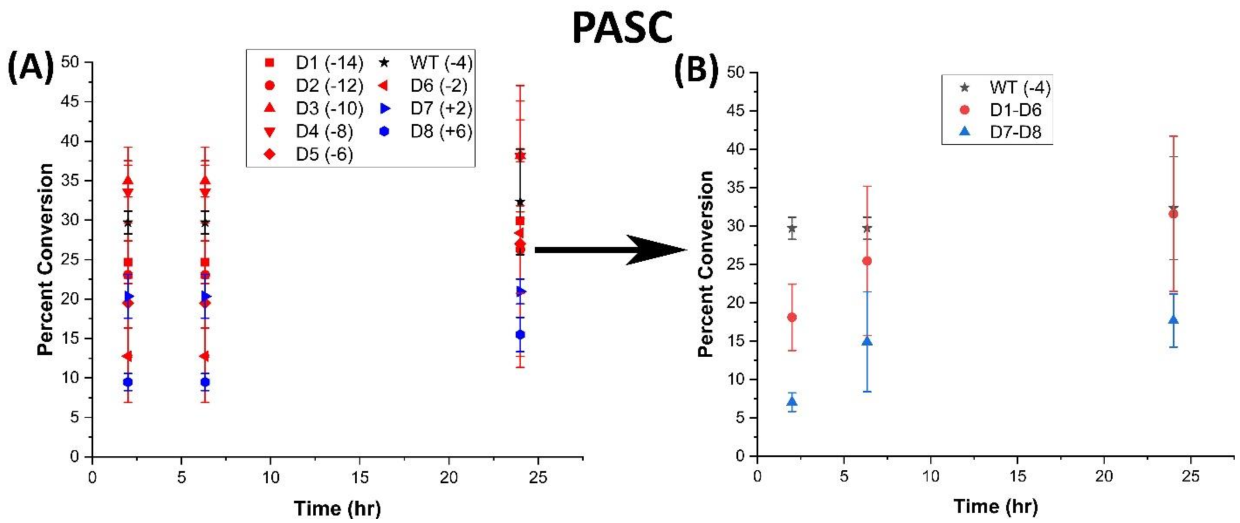


**Figure S3: Hydrolysis of phosphoric acid swollen cellulose (PASC) by CBM2a-Cel5A fusion constructs.** 40 μL of 10 g/L phosphoric acid swollen cellulose (PASC) was hydrolyzed using an enzyme loading of 120 nmol CBM2a Cel5A fusion enzyme per gram cellulose substrate supplemented with 12 nmol of β-glucosidase enzyme (10% of cellulase loading) per gram cellulose substrate for reaction times of 2hrs, 6hrs, and 24hrs. The solubilized reducing sugar concentrations in the supernatant after hydrolysis were determined by the DNS assay method. Percent conversion of biomass to glucose equivalents is reported on the Y-axis. (A) Percent conversion as a function of time (2 hr, 6 hr and 24 hr) for the hydrolysis of PASC by D1 – D8 CBM2a Cel5A and WT CBM2a Cel5A. Error bars represent standard deviation from the mean, based on at least 4 replicates. (B) Based on the data reported in (A), CBM2a Cel5A fusion constructs with negatively charged CBMs (D1 – D6) were grouped together and average hydrolysis yields were obtained for the group, with the error bars representing standard deviation from the mean. Similarly, CBM2a-Cel5A fusion constructs with positively charged CBMs (D7 and D8) were grouped together and average hydrolysis yields were obtained. Trend curves have been added to represent the kinetic profiles of the hydrolysis reaction.

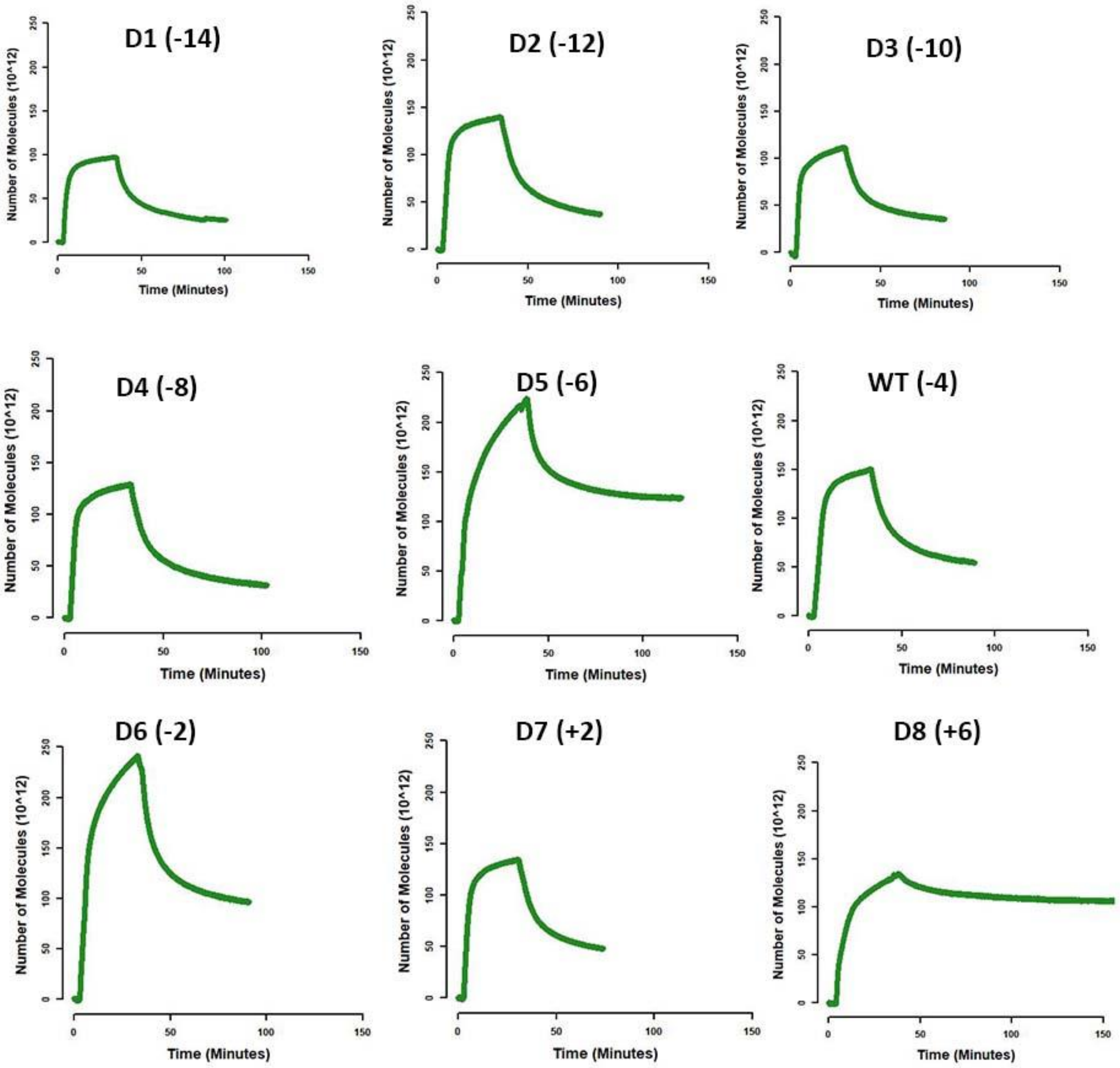


**Figure S4: Binding-unbinding QCM-D representative sensorgrams for GFP-CBM2a designs binding to crystalline cellulose I.** Sensorgrams obtained from Quartz crystal microbalance with dissipation (QCM-D) for binding of GFP-CBM2a designs to cellulose-I were converted to number of molecules * ($10^{12}$) vs time (minutes) using the Sauerbrey equation. These curves were then used to obtain the kinetic parameters for binding and unbinding as reported in **Table 2** of the main manuscript. The equations used for curve fitting have been reported previously in *Nemmaru et al. (2020).*

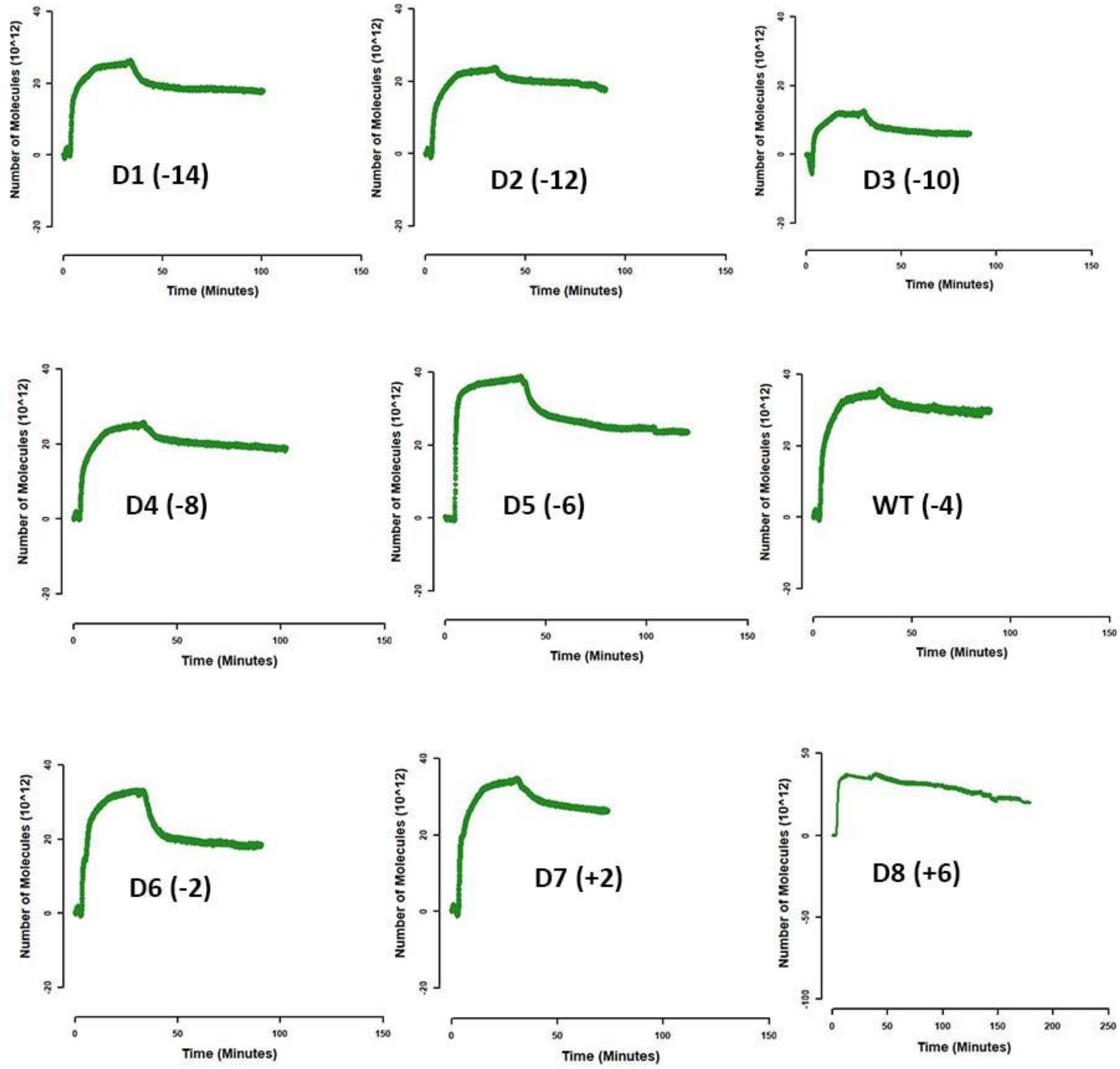


**Figure S5: Binding-unbinding QCM-D representative sensorgrams for GFP-CBM2a designs binding to crystalline cellulose I.** Sensorgrams obtained from Quartz crystal microbalance with dissipation (QCM-D) for binding of GFP-CBM2a designs to lignin were converted to number of molecules * ($10^{12}$) vs time (minutes) using the Sauerbrey equation. These curves were then used to obtain the kinetic parameters for binding and unbinding as reported in **Table 2** of the main manuscript.

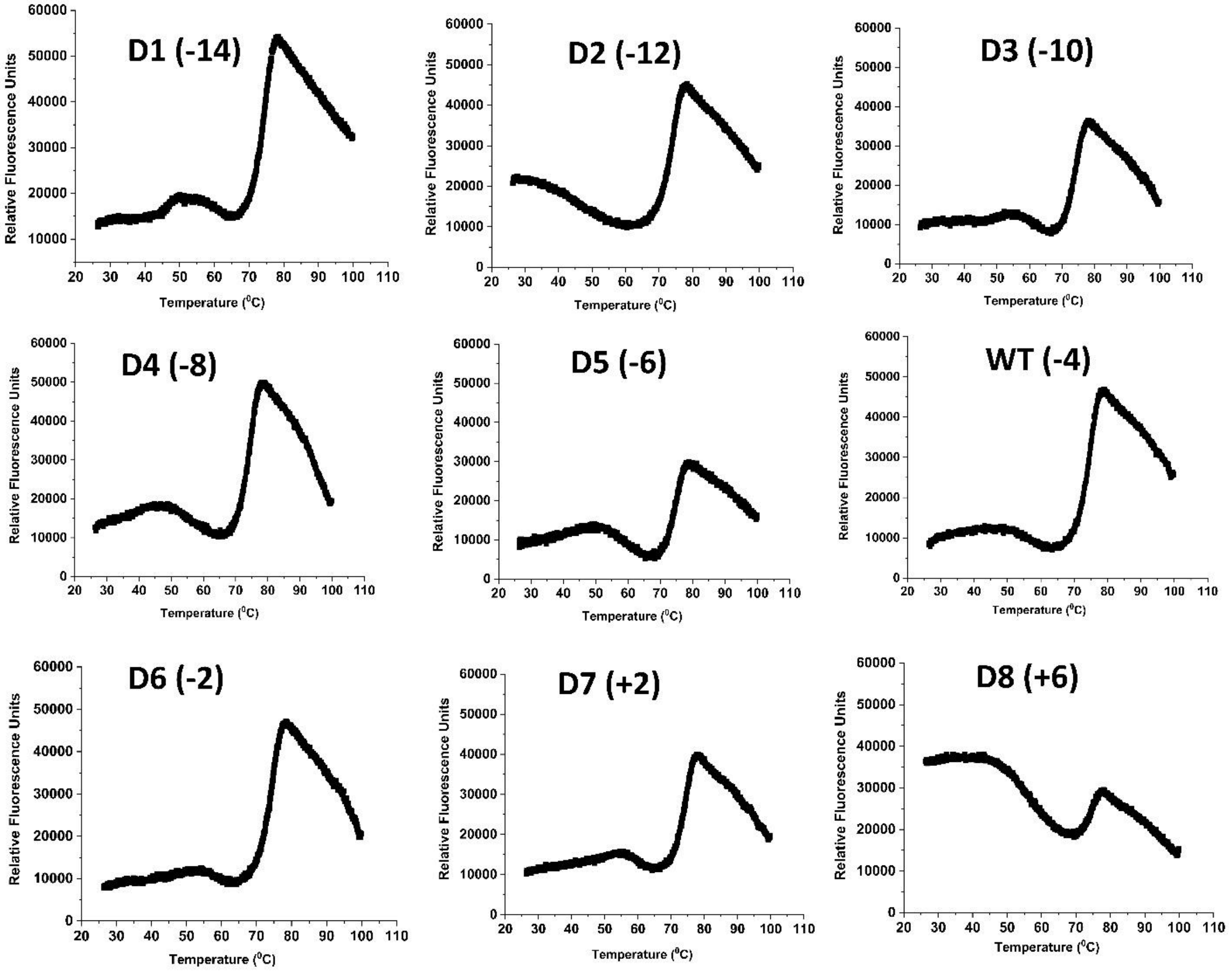


**Figure S6: Representative melt curves from thermal shift assay** 5 µl 200X SYPRO reagent, 5 µl 0.5 M sodium acetate buffer (pH 5.5), enzyme dilution to make up an effective concentration of 5 µM and deionized water to make up the total volume to 50 µl were added to MicroAmp™ EnduraPlate™ 96-well clear microplate (Applied Biosystems$^{TM}$) and fluorescence (excitation – 470 nm; emission – 520 nm) was measured using QuantStudio3 under the FAM dye channel. Representative curves for each CBM2a-Cel5A fusion construct are reported here, starting from D1 (-14 net charge) up to D8 (+6 net charge) including the wild-type (-4 net charge).
