## Supplementary material for "Supercharged Cellulases Show Reduced Non-Productive Binding, But Enhanced Activity, on Pretreated Lignocellulosic Biomass": SI Appendix Sequences

This file contains the following nucleotide and amino acid sequences:

1. Nucleotide sequence for pEC-CBM2a-Cel5A
2. Nucleotide sequence for pEC-GFP-CBM2a
3. Amino acid sequence for wild-type CBM2a with mutable residues highlighted in red

Segments of the open reading frame are highlighted as follows: 8X Histidine Tag, CBM2a, Linker, Cel5A

**> pEC-CBM2a-Cel5A**

AAATAGGGGTTCCGCGCACATTTCCCCGAAAAGTGCCACCTGACGTCTAAGAAACCATTATTATCATGACATTAACCTATAAAAATAGGCGTATCACGAGGCCCTTTCGTCTTCACCTCGAGAAATCATAAAAAATTTATTTGCTTTGTGAGCGGATAACAATTATAATAGATTCAATTGTGAGCGGATAACAATTTCACACAGAATTCATTAAAGAGGAGAAATTAACCATGGGACATCACCATCATCACCATCACCATGGTCTCACCGCCACAGTCACCAAAGAATCCTCGTGGGACAACGGCTACTCCGCGTCCGTCACCGTCCGCAACGACACCTCGAGCACCGTCTCCCAGTGGGAGGTCGTCCTCACCCTGCCCGGCGGCACTACAGTGGCCCAGGTGTGGAACGCCCAGCACACCAGCAGCGGCAACTCCCACACCTTCACCGGGGTTTCCTGGAACAGCACCATCCCGCCCGGAGGCACCGCCTCCTTCGGCTTCATCGCTTCCGGCAGCGGCGAACCCACCCACTGCACCATCAACGGCGCCCCCTGCGACGAAGGCTCCGAGCCGGGCGGCCCCGGCGGTCCCGGAACCCCCTCCCCCGACCCCGGCACGCAGCCCGGCACCGGCACCCCGGTCGAGCGGTACGGCAAAGTCCAGGTCTGCGGCACCCAGCTCTGCGACGAGCACGGCAACCCGGTCCAACTGCGCGGCATGAGCACCCACGGCATCCAGTGGTTCGACCACTGCCTGACCGACAGCTCGCTGGACGCCCTGGCCTACGACTGGAAGGCCGACATCATCCGCCTGTCCATGTACATCCAGGAAGACGGCTACGAGACCAACCCGCGCGGCTTCACCGACCGGATGCACCAGCTCATCGACATGGCCACGGCGCGCGGCCTGTACGTGATCGTGGACTGGCACATCCTCACCCCGGGCGATCCCCACTACAACCTGGACCGGGCCAAGACCTTCTTCGCGGAAATCGCCCAGCGCCACGCCAGCAAGACCAACGTGCTCTACGAGATCGCCAACGAACCCAACGGAGTGAGCTGGGCCTCCATCAAGAGCTACGCCGAAGAGGTCATCCCGGTGATCCGCCAGCGCGACCCCGACTCGGTGATCATCGTGGGCACCCGCGGCTGGTCGTCGCTCGGCGTCTCCGAAGGCTCCGGCCCCGCCGAGATCGCGGCCAACCCGGTCAACGCCTCCAACATCATGTACGCCTTCCACTTCTACGCGGCCTCGCACCGCGACAACTACCTCAACGCGCTGCGTGAGGCCTCCGAGCTGTTCCCGGTCTTCGTCACCGAGTTCGGCACCGAGACCTACACCGGTGACGGCGCCAACGACTTCCAGATGGCCGACCGCTACATCGACCTGATGGCGGAACGGAAGATCGGGTGGACCAAGTGGAACTACTCGGACGACTTCCGTTCCGGCGCGGTCTTCCAGCCGGGCACCTGCGCGTCCGGCGGCCCGTGGAGCGGTTCGTCGCTGAAGGCGTCCGGACAGTGGGTGCGGAGCAAGCTCCAGTCCTAGCCCAAACGAATTCGAGCTCGGTACCCGGGGATCCTCTAGAGTCGACCTGCAGGCATGCAAGCTGATCCGGCTGCTAACAAAGCCCGAAAGGAAGCTGAGTTGGCTGCTGCCACCGCTGAGCAATAACTAGCAAACTCGTTTCTCGTTCAGCTTTCTTGTACAAAGTGGTGATGGCGCGCCTGTAGGACGTCGACGGTACCATCGATACGCGTTCGAAGCTTAATTAGCTGAGCTTGGACTCCTGTTGATAGATCCAGTAATGACCTCAGAACTCCATCTGGATTTGTTCAGAACGCTCGGTTGCCGCCGGGCGTTTTTTATTGGTGAGAATCCAAGCTAGCTTGGCGAGATTTTCAGGAGCTAAGGAAGCTAAAATGGAGAAAAAAATCACTGGATATACCACCGTTGATATATCCCAATGGCATCGTAAAGAACATTTTGAGGCATTTCAGTCAGTTGCTCAATGTACCTATAACCAGACCGTTCAGCTGGATATTACGGCCTTTTTAAAGACCGTAAAGAAAAATAAGCACAAGTTTTATCCGGCCTTTATTCACATTCTTGCCCGCCTGATGAATGCTCATCCGGAATTTCGTATGGCAATGAAAGACGGTGAGCTGGTGATATGGGATAGTGTTCACCCTTGTTACACCGTTTTCCATGAGCAAACTGAAACGTTTTCATCGCTCTGGAGTGAATACCACGACGATTTCCGGCAGTTTCTACACATATATTCGCAAGATGTGGCGTGTTACGGTGAAAACCTGGCCTATTTCCCTAAAGGGTTTATTGAGAATATGTTTTTCGTCTCAGCCAATCCCTGGGTGAGTTTCACCAGTTTTGATTTAAACGTGGCCAATATGGACAACTTCTTCGCCCCCGTTTTCACCATGGGCAAATATTATACGCAAGGCGACAAGGTGCTGATGCCGCTGGCGATTCAGGTTCATCATGCCGTTTGTGATGGCTTCCATGTCGGCAGAATGCTTAATGAATTACAACAGTACTGCGATGAGTGGCAGGGCGGGGCGTAATTTTTTTAAGGCAGTTATTGGTGCCCTTAAACGCCTGGGGTAATGACTCTCTAGCTTGAGGCATCAAATAAAACGAAAGGCTCAGTCGAAAGACTGGGCCTTTCGTTTTATCTGTTGTTTGTCGGTGAACGCTCTCCTGAGTAGGACAAATCCGCCCTCTAGATTACGTGCAGTCGATGATAAGCTGTCAAACATGAGAATTGTGCCTAATGAGTGAGCTAACTTACATTAATTGCGTTGCGCTCACTGCCCGCTTTCCAGTCGGGAAACCTGTCGTGCCAGCTGCATTAATGAATCGGCCAACGCGCGGGGAGAGGCGGTTTGCGTATTGGGCGCCAGGGTGGTTTTTCTTTTCACCAGTGAGACGGGCAACAGCTGATTGCCCTTCACCGCCTGGCCCTGAGAGAGTTGCAGCAAGCGGTCCACGCTGGTTTGCCCCAGCAGGCGAAAATCCTGTTTGATGGTGGTTAACGGCGGGATATAACATGAGCTGTCTTCGGTATCGTCGTATCCCACTACCGAGATATCCGCACCAACGCGCAGCCCGGACTCGGTAATGGCGCGCATTGCGCCCAGCGCCATCTGATCGTTGGCAACCAGCATCGCAGTGGGAACGATGCCCTCATTCAGCATTTGCATGGTTTGTTGAAAACCGGACATGGCACTCCAGTCGCCTTCCCGTTCCGCTATCGGCTGAATTTGATTGCGAGTGAGATATTTATGCCAGCCAGCCAGACGCAGACGCGCCGAGACAGAACTTAATGGGCCCGCTAACAGCGCGATTTGCTGGTGACCCAATGCGACCAGATGCTCCACGCCCAGTCGCGTACCGTCTTCATGGGAGAAAATAATACTGTTGATGGGTGTCTGGTCAGAGACATCAAGAAATAACGCCGGAACATTAGTGCAGGCAGCTTCCACAGCAATGGCATCCTGGTCATCCAGCGGATAGTTAATGATCAGCCCACTGACGCGTTGCGCGAGAAGATTGTGCACCGCCGCTTTACAGGCTTCGACGCCGCTTCGTTCTACCATCGACACCACCACGCTGGCACCCAGTTGATCGGCGCGAGATTTAATCGCCGCGACAATTTGCGACGGCGCGTGCAGGGCCAGACTGGAGGTGGCAACGCCAATCAGCAACGACTGTTTGCCCGCCAGTTGTTGTGCCACGCGGTTGGGAATGTAATTCAGCTCCGCCATCGCCGCTTCCACTTTTTCCCGCGTTTTCGCAGAAACGTGGCTGGCCTGGTTCACCACGCGGGAAACGGTCTGATAAGAGACACCGGCATACTCTGCGACATCGTATAACGTTACTGGTTTCACATTCACCACCCTGAATTGACTCTCTTCCGGGCGCTATCATGCCATACCGCGAAAGGTTTTGCACCATTCGATGGTGTCGGAATTTCGGGCAGCGTTGGGTCCTGGCCACGGGTGCGCATGATCTAGAGCTGCCTCGCGCGTTTCGGTGATGACGGTGAAAACCTCTGACACATGCAGCTCCCGGAGACGGTCACAGCTTGTCTGTAAGCGGATGCCGGGAGCAGACAAGCCCGTCAGGGCGCGTCAGCGGGTGTTGGCGGGTGTCGGGGCGCAGCCATGACCCAGTCACGTAGCGATAGCGGAGTGTATACTGGCTTAACTATGCGGCATCAGAGCAGATTGTACTGAGAGTGCACCATATGCGGTGTGAAATACCGCACAGATGCGTAAGGAGAAAATACCGCATCAGGCGCTCTTCCGCTTCCTCGCTCACTGACTCGCTGCGCTCGGTCGTTCGGCTGCGGCGAGCGGTATCAGCTCACTCAAAGGCGGTAATACGGTTATCCACAGAATCAGGGGATAACGCAGGAAAGAACATGTGAGCAAAAGGCCAGCAAAAGGCCAGGAACCGTAAAAAGGCCGCGTTGCTGGCGTTTTTCCATAGGCTCCGCCCCCCTGACGAGCATCACAAAAATCGACGCTCAAGTCAGAGGTGGCGAAACCCGACAGGACTATAAAGATACCAGGCGTTTCCCCCTGGAAGCTCCCTCGTGCGCTCTCCTGTTCCGACCCTGCCGCTTACCGGATACCTGTCCGCCTTTCTCCCTTCGGGAAGCGTGGCGCTTTCTCATAGCTCACGCTGTAGGTATCTCAGTTCGGTGTAGGTCGTTCGCTCCAAGCTGGGCTGTGTGCACGAACCCCCCGTTCAGCCCGACCGCTGCGCCTTATCCGGTAACTATCGTCTTGAGTCCAACCCGGTAAGACACGACTTATCGCCACTGGCAGCAGCCACTGGTAACAGGATTAGCAGAGCGAGGTATGTAGGCGGTGCTACAGAGTTCTTGAAGTGGTGGCCTAACTACGGCTACACTAGAAGGACAGTATTTGGTATCTGCGCTCTGCTGAAGCCAGTTACCTTCGGAAAAAGAGTTGGTAGCTCTTGATCCGGCAAACAAACCACCGCTGGTAGCGGTGGTTTTTTTGTTTGCAAGCAGCAGATTACGCGCAGAAAAAAAGGATCTCAAGAAGATCCTTTGATCTTTTCTACGGGGTCTGACGCTCAGTGGAACGAAAACTCACGTTAAGGGATTTTGGTCATGAGATTATCAAAAAGGATCTTCACCTAGATCCTTTTAAATTAAAAATGAAGTTTTAAATCAATCTAAAGTATATATGAGTAAACTTGGTCTGACAGCCTAGGTCAGAAGAACTCGTCAAGAAGGCGATAGAAGGCGATGCGCTGCGAATCGGGAGCGGCGATACCGTAAAGCACGAGGAAGCGGTCAGCCCATTCGCCGCCAAGCTCTTCAGCAATATCACGGGTAGCCAACGCTATGTCCTGATAGCGGTCCGCCACACCCAGCCGGCCACAGTCGATGAATCCAGAAAAGCGGCCATTTTCCACCATGATATTCGGCAAGCAGGCATCGCCATGAGTCACGACGAGATCCTCACCATCCGGCATACGCGCCTTGAGCCTGGCGAACAGTTCGGCTGGCGCGAGCCCCTGATGCTCTTCGTCCAGATCATCCTGATCGACAAGACCGGCTTCCATCCGAGTGCGTGCTCGCTCGATGCGATGTTTCGCTTGGTGGTCGAATGGGCAGGTAGCCGGATCAAGCGTATGCAGCCGCCGCATTGCATCAGCCATGATGGATACTTTCTCGGCAGGAGCAAGGTGAGATGACAGGAGATCCTGCCCCGGCACTTCGCCCAATAGCAGCCAGTCCCTTCCCGCTTCAGTGACAACGTCGAGCACAGCTGCGCAAGGAACGCCCGTCGTGGCCAGCCACGATAGCCGCGCTGCCTCGTCCTGCAGTTCATTCAGGGCACCGGACAGGTCGGTCTTGACAAAAAGAACCGGGCGCCCCTGCGCTGACAGCCGGAACACGGCGGCATCAGAGCAGCCGATTGTCTGTTGTGCCCAGTCATAGCCGAATAGCCTCTCCACCCAAGCGGCCGGAGAACCTGCGTGCAATCCATCTTGTTCAAGCATGCGAAACGACCGTCATCCTGTCTCTTGATCAGATCTTGATCCCCTGCGCCATCAGATCCTTGGCGGCAAGAAAGCCATCCAGTTTACTTTGCAGGGCTTCCCAACCTTACCAGAGGGCGCCCCAGCTGGCAATTCTTTTGAAGCTCACGCTGCCGCAAGCACTCAGGGCGTACG

Segments of the open reading frame are highlighted as follows: 8X Histidine Tag, GFP, Linker, CBM2a

> **pEC-GFP-CBM2a**

AAATAGGGGTTCCGCGCACATTTCCCCGAAAAGTGCCACCTGACGTCTAAGAAACCATTATTATCATGACATTAACCTATAAAAATAGGCGTATCACGAGGCCCTTTCGTCTTCACCTCGAGAAATCATAAAAAATTTATTTGCTTTGTGAGCGGATAACAATTATAATAGATTCAATTGTGAGCGGATAACAATTTCACACAGAATTCATTAAAGAGGAGAAATTAACCATGGGACATCACCATCATCACCATCACCATGCATCCGAAAACCTGTACTTCCAGGCGATCGCCTCCAAAGGTGAAGAACTGTTCACCGGTGTTGTTCCGATCCTGGTTGAACTGGACGGTGACGTTAACGGTCACAAATTCTCCGTTTCCGGTGAAGGTGAAGGTGACGCTACCTACGGTAAACTGACCCTGAAATTCATCTGCACCACCGGTAAACTGCCGGTTCCGTGGCCGACCCTGGTTACCACCCTGACCTACGGTGTTCAGTGCTTCTCCCGTTACCCGGACCACATGAAACAGCACGACTTCTTCAAATCCGCTATGCCGGAAGGTTACGTTCAGGAACGTACCATCTCCTTCAAAGACGACGGTAACTACAAAACCCGTGCTGAAGTTAAATTCGAAGGTGACACCCTGGTTAACCGTATCGAACTGAAAGGTATCGACTTCAAAGAAGACGGTAACATCCTGGGTCACAAACTGGAATACAACTACAACTCCCACAACGTTTACATCACCGCTGACAAACAGAAAAACGGTATCAAAGCTAACTTCAAAATCCGTCACAACATCGAAGACGGTTCCGTTCAGCTGGCTGACCACTACCAGCAGAACACCCCGATCGGTGACGGTCCGGTTCTGCTGCCGGACAACCACTACCTGTCCACCCAGTCCGCTCTGTCCAAAGACCCGAACGAAAAACGTGACCACATGGTTCTGCTGGAATTCGTTACCGCTGCTGGTATCACCCACGGTATGGACGAACTGTACAAAGGTTTAAACGCGACTCCCACTAAAGGTGCCACTCCTACCAATACGGCGACTCCGACTAAGTCGGCAACGGCAACGCCCACTCGCCCCAGCGTACCGACCAATACTCCGACTAATACCCCGGCGAACACCCTTAAGGCCGGCTGCTCGGTGGACTACACGGTCAACTCCTGGGGTACCGGGTTCACCGCCAACGTCACCATCACCAACCTCGGCAGTGCGATCAACGGCTGGACCCTGGAGTGGGACTTCCCCGGCAACCAGCAGGTGACCAACCTGTGGAACGGGACCTACACCCAGTCCGGGCAGCACGTGTCGGTCAGCAACGCCCCGTACAACGCCTCCATCCCGGCCAACGGAACGGTTGAGTTCGGGTTCAACGGCTCCTACTCGGGCAGCAACGACATCCCCTCCTCCTTCAAGCTGAACGGGGTTACCTGCTAAGGATCCTCTAGAGTCGACCTGCAGGCATGCAAGCTGATCCGGCTGCTAACAAAGCCCGAAAGGAAGCTGAGTTGGCTGCTGCCACCGCTGAGCAATAACTAGCAAACTCGTTTCTCGTTCAGCTTTCTTGTACAAAGTGGTGATGGCGCGCCTGTAGGACGTCGACGGTACCATCGATACGCGTTCGAAGCTTAATTAGCTGAGCTTGGACTCCTGTTGATAGATCCAGTAATGACCTCAGAACTCCATCTGGATTTGTTCAGAACGCTCGGTTGCCGCCGGGCGTTTTTTATTGGTGAGAATCCAAGCTAGCTTGGCGAGATTTTCAGGAGCTAAGGAAGCTAAAATGGAGAAAAAAATCACTGGATATACCACCGTTGATATATCCCAATGGCATCGTAAAGAACATTTTGAGGCATTTCAGTCAGTTGCTCAATGTACCTATAACCAGACCGTTCAGCTGGATATTACGGCCTTTTTAAAGACCGTAAAGAAAAATAAGCACAAGTTTTATCCGGCCTTTATTCACATTCTTGCCCGCCTGATGAATGCTCATCCGGAATTTCGTATGGCAATGAAAGACGGTGAGCTGGTGATATGGGATAGTGTTCACCCTTGTTACACCGTTTTCCATGAGCAAACTGAAACGTTTTCATCGCTCTGGAGTGAATACCACGACGATTTCCGGCAGTTTCTACACATATATTCGCAAGATGTGGCGTGTTACGGTGAAAACCTGGCCTATTTCCCTAAAGGGTTTATTGAGAATATGTTTTTCGTCTCAGCCAATCCCTGGGTGAGTTTCACCAGTTTTGATTTAAACGTGGCCAATATGGACAACTTCTTCGCCCCCGTTTTCACCATGGGCAAATATTATACGCAAGGCGACAAGGTGCTGATGCCGCTGGCGATTCAGGTTCATCATGCCGTTTGTGATGGCTTCCATGTCGGCAGAATGCTTAATGAATTACAACAGTACTGCGATGAGTGGCAGGGCGGGGCGTAATTTTTTTAAGGCAGTTATTGGTGCCCTTAAACGCCTGGGGTAATGACTCTCTAGCTTGAGGCATCAAATAAAACGAAAGGCTCAGTCGAAAGACTGGGCCTTTCGTTTTATCTGTTGTTTGTCGGTGAACGCTCTCCTGAGTAGGACAAATCCGCCCTCTAGATTACGTGCAGTCGATGATAAGCTGTCAAACATGAGAATTGTGCCTAATGAGTGAGCTAACTTACATTAATTGCGTTGCGCTCACTGCCCGCTTTCCAGTCGGGAAACCTGTCGTGCCAGCTGCATTAATGAATCGGCCAACGCGCGGGGAGAGGCGGTTTGCGTATTGGGCGCCAGGGTGGTTTTTCTTTTCACCAGTGAGACGGGCAACAGCTGATTGCCCTTCACCGCCTGGCCCTGAGAGAGTTGCAGCAAGCGGTCCACGCTGGTTTGCCCCAGCAGGCGAAAATCCTGTTTGATGGTGGTTAACGGCGGGATATAACATGAGCTGTCTTCGGTATCGTCGTATCCCACTACCGAGATATCCGCACCAACGCGCAGCCCGGACTCGGTAATGGCGCGCATTGCGCCCAGCGCCATCTGATCGTTGGCAACCAGCATCGCAGTGGGAACGATGCCCTCATTCAGCATTTGCATGGTTTGTTGAAAACCGGACATGGCACTCCAGTCGCCTTCCCGTTCCGCTATCGGCTGAATTTGATTGCGAGTGAGATATTTATGCCAGCCAGCCAGACGCAGACGCGCCGAGACAGAACTTAATGGGCCCGCTAACAGCGCGATTTGCTGGTGACCCAATGCGACCAGATGCTCCACGCCCAGTCGCGTACCGTCTTCATGGGAGAAAATAATACTGTTGATGGGTGTCTGGTCAGAGACATCAAGAAATAACGCCGGAACATTAGTGCAGGCAGCTTCCACAGCAATGGCATCCTGGTCATCCAGCGGATAGTTAATGATCAGCCCACTGACGCGTTGCGCGAGAAGATTGTGCACCGCCGCTTTACAGGCTTCGACGCCGCTTCGTTCTACCATCGACACCACCACGCTGGCACCCAGTTGATCGGCGCGAGATTTAATCGCCGCGACAATTTGCGACGGCGCGTGCAGGGCCAGACTGGAGGTGGCAACGCCAATCAGCAACGACTGTTTGCCCGCCAGTTGTTGTGCCACGCGGTTGGGAATGTAATTCAGCTCCGCCATCGCCGCTTCCACTTTTTCCCGCGTTTTCGCAGAAACGTGGCTGGCCTGGTTCACCACGCGGGAAACGGTCTGATAAGAGACACCGGCATACTCTGCGACATCGTATAACGTTACTGGTTTCACATTCACCACCCTGAATTGACTCTCTTCCGGGCGCTATCATGCCATACCGCGAAAGGTTTTGCACCATTCGATGGTGTCGGAATTTCGGGCAGCGTTGGGTCCTGGCCACGGGTGCGCATGATCTAGAGCTGCCTCGCGCGTTTCGGTGATGACGGTGAAAACCTCTGACACATGCAGCTCCCGGAGACGGTCACAGCTTGTCTGTAAGCGGATGCCGGGAGCAGACAAGCCCGTCAGGGCGCGTCAGCGGGTGTTGGCGGGTGTCGGGGCGCAGCCATGACCCAGTCACGTAGCGATAGCGGAGTGTATACTGGCTTAACTATGCGGCATCAGAGCAGATTGTACTGAGAGTGCACCATATGCGGTGTGAAATACCGCACAGATGCGTAAGGAGAAAATACCGCATCAGGCGCTCTTCCGCTTCCTCGCTCACTGACTCGCTGCGCTCGGTCGTTCGGCTGCGGCGAGCGGTATCAGCTCACTCAAAGGCGGTAATACGGTTATCCACAGAATCAGGGGATAACGCAGGAAAGAACATGTGAGCAAAAGGCCAGCAAAAGGCCAGGAACCGTAAAAAGGCCGCGTTGCTGGCGTTTTTCCATAGGCTCCGCCCCCCTGACGAGCATCACAAAAATCGACGCTCAAGTCAGAGGTGGCGAAACCCGACAGGACTATAAAGATACCAGGCGTTTCCCCCTGGAAGCTCCCTCGTGCGCTCTCCTGTTCCGACCCTGCCGCTTACCGGATACCTGTCCGCCTTTCTCCCTTCGGGAAGCGTGGCGCTTTCTCATAGCTCACGCTGTAGGTATCTCAGTTCGGTGTAGGTCGTTCGCTCCAAGCTGGGCTGTGTGCACGAACCCCCCGTTCAGCCCGACCGCTGCGCCTTATCCGGTAACTATCGTCTTGAGTCCAACCCGGTAAGACACGACTTATCGCCACTGGCAGCAGCCACTGGTAACAGGATTAGCAGAGCGAGGTATGTAGGCGGTGCTACAGAGTTCTTGAAGTGGTGGCCTAACTACGGCTACACTAGAAGGACAGTATTTGGTATCTGCGCTCTGCTGAAGCCAGTTACCTTCGGAAAAAGAGTTGGTAGCTCTTGATCCGGCAAACAAACCACCGCTGGTAGCGGTGGTTTTTTTGTTTGCAAGCAGCAGATTACGCGCAGAAAAAAAGGATCTCAAGAAGATCCTTTGATCTTTTCTACGGGGTCTGACGCTCAGTGGAACGAAAACTCACGTTAAGGGATTTTGGTCATGAGATTATCAAAAAGGATCTTCACCTAGATCCTTTTAAATTAAAAATGAAGTTTTAAATCAATCTAAAGTATATATGAGTAAACTTGGTCTGACAGCCTAGGTCAGAAGAACTCGTCAAGAAGGCGATAGAAGGCGATGCGCTGCGAATCGGGAGCGGCGATACCGTAAAGCACGAGGAAGCGGTCAGCCCATTCGCCGCCAAGCTCTTCAGCAATATCACGGGTAGCCAACGCTATGTCCTGATAGCGGTCCGCCACACCCAGCCGGCCACAGTCGATGAATCCAGAAAAGCGGCCATTTTCCACCATGATATTCGGCAAGCAGGCATCGCCATGAGTCACGACGAGATCCTCACCATCCGGCATACGCGCCTTGAGCCTGGCGAACAGTTCGGCTGGCGCGAGCCCCTGATGCTCTTCGTCCAGATCATCCTGATCGACAAGACCGGCTTCCATCCGAGTGCGTGCTCGCTCGATGCGATGTTTCGCTTGGTGGTCGAATGGGCAGGTAGCCGGATCAAGCGTATGCAGCCGCCGCATTGCATCAGCCATGATGGATACTTTCTCGGCAGGAGCAAGGTGAGATGACAGGAGATCCTGCCCCGGCACTTCGCCCAATAGCAGCCAGTCCCTTCCCGCTTCAGTGACAACGTCGAGCACAGCTGCGCAAGGAACGCCCGTCGTGGCCAGCCACGATAGCCGCGCTGCCTCGTCCTGCAGTTCATTCAGGGCACCGGACAGGTCGGTCTTGACAAAAAGAACCGGGCGCCCCTGCGCTGACAGCCGGAACACGGCGGCATCAGAGCAGCCGATTGTCTGTTGTGCCCAGTCATAGCCGAATAGCCTCTCCACCCAAGCGGCCGGAGAACCTGCGTGCAATCCATCTTGTTCAAGCATGCGAAACGACCGTCATCCTGTCTCTTGATCAGATCTTGATCCCCTGCGCCATCAGATCCTTGGCGGCAAGAAAGCCATCCAGTTTACTTTGCAGGGCTTCCCAACCTTACCAGAGGGCGCCCCAGCTGGCAATTCTTTTGAAGCTCACGCTGCCGCAAGCACTCAGGGCGTACG

> CBM2a wild-type

AGCSVDYTVNSWGTGFTANVTITNLG**S**AI**N**GW**T**LEWDFPG**N**QQVTNLWNG**T**Y**T**Q**S**G**Q**HVSV**S**NAPYNASIPANGTVEFGFNG**S**Y**S**GSNDIPSSFKLNGVTC

1. ^#^ Equal contribution
